# Arena3D 3.0: a web server for exploring, analyzing, annotating and sharing multilayered networks in 3D

**DOI:** 10.64898/2026.10.23.761450

**Authors:** Evangelos Karatzas, Fotis Baltoumas, Maria Kokoli, Alexandros Galaras, Eleni Aplakidou, Marios Itsios, Iro N. Chasapi, Maria N. Chasapi, Efthimia Lamari, Ilias Georgakopoulos-Soares, Nataša Pržulj, Georgios A. Pavlopoulos

## Abstract

Multilayer networks describe biological entities linked by several kinds of relation or measured under several conditions. Arena3D 3.0, a complete reimplementation of Arena3D^web^, gives each layer its own place in an interactive 3D scene. Networks are uploaded or imported from STRING, IntAct, OmniPath, NDEx, Reactome and the Gene Ontology. Twelve layouts, seven in 3D, are available, and node copies can be aligned across layers. Layers are drawn as floors or 3D containers in thirteen arrangements, including 3D hive plots. Communities and annotated groups can be laid out apart and outlined as hulls or fog clouds, and edges can be bundled. Analyses cover node and multilayer metrics, distributions and perturbation, with chord, Sankey, heatmap, UpSet and UMAP views. Networks go to Cytoscape, Gephi Lite, NORMA and JBrowse 2, gene lists to Flame, and a REST API gives programmatic access. Arena3D 3.0 is free at www.arena3d.org without login.

**GRAPHICAL ABSTRACT:** 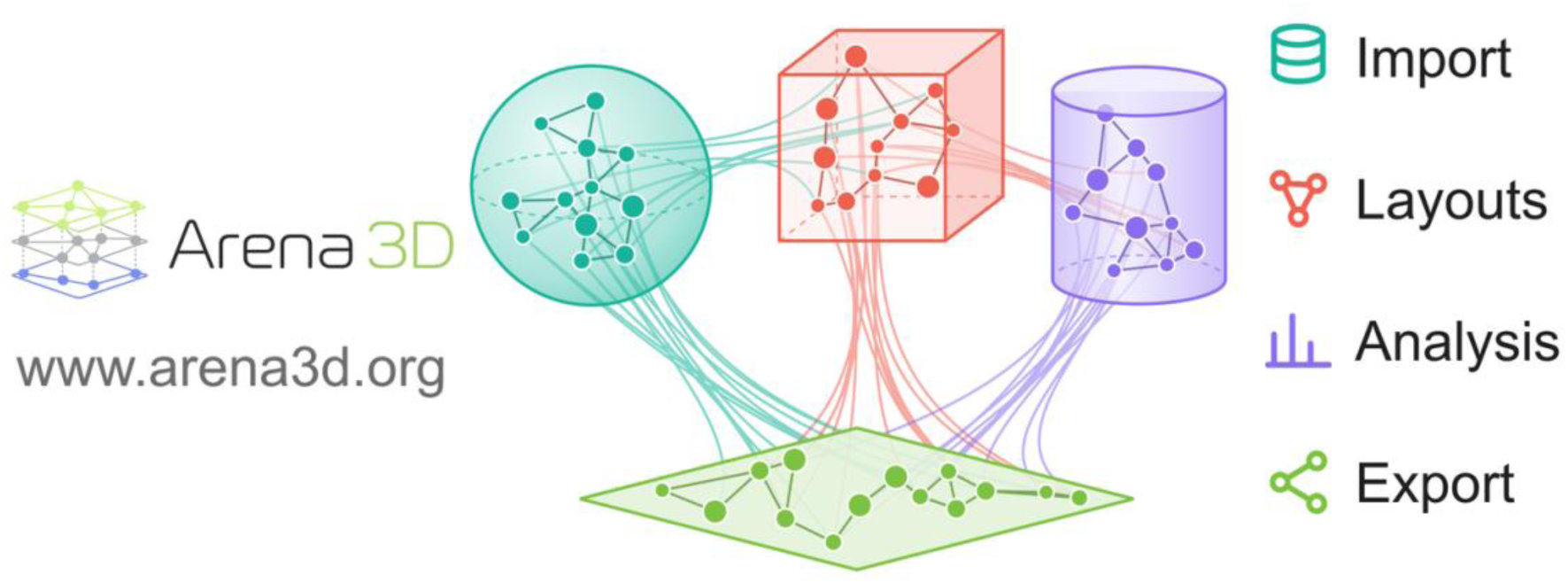

## INTRODUCTION

A protein may bind some partners, regulate others, be coexpressed with a third group and share pathways with a fourth, and each of these relations comes from a different kind of evidence (1, 2). Multilayer networks keep such relations apart (3). Each kind of relation forms a layer, and links between layers connect the copies of the same entity or related entities (4, 5). The same structure fits a network measured across tissues, time points or conditions, or interactions collected from several databases, and bipartite and multipartite graphs are special cases of it (6).

Most network viewers either flatten multilayer data into a single graph or place the layers side by side (7–15). General-purpose tools such as Cytoscape (16), Gephi (17), VisANT (18), NAViGaTOR (19) and Cytoscape.js (20), as well as the igraph (21) and NetworkX (22) libraries, store layers as node or edge attributes. Viewers built for 3D and immersive settings, such as BioLayout Express3D and its successor Graphia (23, 24), iCAVE (25) and VRNetzer (26), place a single graph in space. Software written for multilayer networks, including MuxViz (27), the R packages multinet (28) and mully (29) and the Python library pymnet (30), focuses on analysis and must be installed locally. Apart from the graphical interface of MuxViz, these tools offer static or scripted views (31).

Arena3D was first released in 2008 as a Java desktop application that stacked biological networks as layers in 3D (32). A later version added the analysis of time-course data through clustering and correlation (33). Arena3D^web^ then moved the concept to the web, with each layer on its own floor and the edges between layers drawn through space (34). Its second release introduced directed edges, up to nine information channels per node pair, clustering, predefined layer arrangements, a virtual reality mode and an API together with a Cytoscape app (35).

Arena3D^web^ could compute one joint layout for selected layers, but it could not balance the alignment of node copies against the structure of each layer. In the two networks benchmarked below, independent layouts leave the copies of a protein 26% and 29% of the layout diagonal apart, and the edges between layers cross one another. Nodes could only sit on flat floors. Clusters and annotations changed the colors of the nodes but not the layout, even though NORMA (36) and NORMA 2.0 (37) had introduced group-aware layouts that keep annotated groups apart. Edges were drawn as straight lines, or as curves when a pair of nodes had several channels and could not be bundled. The analysis was limited to a few node metrics, and the 3D scene was the only available view.

Arena3D 3.0 addresses these points (Table 1). It keeps the visual concept and the file formats of the earlier versions and adds alignment across layers, 3D containers, a hive plot (38) in 3D, group layouts, edge bundling, multilayer analysis, alternative views such as UpSet plots (39) and a documented REST API.

**Table 1.** Features of Arena3D 3.0 compared with the earlier releases.

| Feature | Arena3D (2008) | Arena3D (2012) | Arena3D<sup>web</sup> (2021) | Arena3D<sup>web</sup> (2023) | Arena3D 3.0 |
| --- | --- | --- | --- | --- | --- |
| **Implementation** | Java desktop application | Java desktop application | R, Shiny and JavaScript web server | R, Shiny and JavaScript web server | FastAPI and igraph service with a TypeScript and three.js client |
| **Availability today** | Discontinued, no longer distributed as a standalone application | Discontinued, no longer distributed as a standalone application | Superseded by Arena3D 3.0 | Superseded by Arena3D 3.0 | Current version at www.arena3d.org and as a Docker image |
| **Runs in a web browser** | No | No | Yes | Yes | Yes, on desktops, tablets and phones |
| **Limits on the public server** | *Not applicable* | *Not applicable* | 10 layers | 10 layers | 20 layers, 10 channels, 20,000 nodes and 10,000 edges by default |
| **Input** | Tab Delimited File | Time-course expression data | Uploaded Tab Delimited files | Sessions, API, Cytoscape app | Arena3D TSV, edge lists, GraphML, GML, SIF and CX2, attribute, group and value files, sessions |
| **Database import** | No | No | No | No | STRING, IntAct, OmniPath, NDEx, Reactome and Gene Ontology |
| **Several networks** | One at a time | One at a time | One at a time | One at a time | Several loaded at once, a Networks menu to switch, rename and delete them, network comparison |
| **Edges and channels** | Weighted, undirected | Weighted, undirected | Weighted, undirected | Directed, up to 9 channels per pair | Directed, up to 10 channels drawn parallel, curved or as one line, channel weights in layouts |
| **Layouts** | 2D, per layer | 2D, per layer | 2D, per layer or layers together | 2D, per layer or layers together | Twelve algorithms, seven also in 3D, per layer, layers together or selected nodes |
| **UMAP layouts and view** | No | No | No | No | By topology, attributes or both, in 2D or 3D, and as a scatter plot view |
| **Alignment across layers** | No | Partially | Joint layout of selected layers | Joint layout of selected layers | Exact, soft, anchor and chain alignment with a misalignment value |
| **Single 3D network mode** | No | No | No | No | Yes, with a Spread slider |
| **Layer arrangements** | Staggered layers in 3D | Staggered layers in 3D | Floors, per-layer transforms | Zigzag, star and cube arrangements | Thirteen, including helix, circle, stack, fan, chord diagram and hive plot in a plane or in 3D |
| **Layer shapes** | Floors | Floors | Floors | Floors | Square or round floors in five styles, or 13 3D containers with nodes inside or on the surface, per-layer axes |
| **Node shapes** | Spheres | Spheres | Spheres | Spheres | Fourteen shapes per node or per layer, borders and proportions |
| **Label styles** | No | No | No | No | Halo by default, box, shadow or plain, four positions, colors by group, chosen labels and a label picker |
| **Community detection** | No | Clustering of time-course data | No | Clustering | Five algorithms including multiplex Leiden, communities saved as annotation sets |
| **Group-aware layouts** | No | No | No | No | NORMA 2.0 *virtual node*, *group gravity* and *super node* strategies on clusters, annotations or layers |
| **Group annotations** | No | No | No | No | Hulls, outlines, clouds, spheres, ellipsoids or boxes, pie charts, group statistics and overlaps |
| **Edge clutter** | No | No | No | Channel curvature | Force-directed edge bundling within and between layers or over the whole graph |
| **Analysis** | No | Correlations of time-course data | Topology-based highlighting | Clustering | Node metrics on undirected, directed or multiedge graphs, network statistics, layer similarity, enrichment |
| Multilayer metrics | No | No | No | No | Multiplex degree, participation coefficient and multiplex PageRank |
| Plots | No | No | No | No | Distributions with power-law fit, correlations, rich-club, k-core shells, centrality rankings, layer profiles |
| Perturbation analysis | No | No | No | No | Node and edge knockouts, network propagation, attack and robustness curves |
| Search | No | No | Basic search | Basic search | Nodes and edges by exact, prefix, suffix, substring or regular expression match, with result tables and camera focus |
| Selection and paths | No | No | No | No | Freehand lasso, components, hubs and layers, live shortest paths among selected nodes |
| Styling and editing | Node color and size | Node color and size | Node color and size | Node color and size | Style and attribute tables modeled on Cytoscape, inspector, attribute files that load back |
| Themes and color vision | No | No | No | Light, dark and gray themes | 27 themes, eight of them color-blind safe, color-blind mode and simulator |
| Phones and tablets | No | No | No | No | Touch gestures and panels as bottom sheets |
| Virtual reality | No | No | No | VR mode | Not yet carried over |
| Alternative views | No | No | No | No | Chord, Sankey, heatmap, interactive UpSet and UMAP views |
| Export formats | Tab Delimited | Tab Delimited | Images | JSON sessions | GraphML, CX2, Cytoscape.js, SIF, MuxViz, PNG, SVG, GLB, video and sessions |
| Transfers to other tools | No | No | No | From the Cytoscape app | Cytoscape, Gephi Lite, Vega editor, STRING, NORMA, Flame, Gosling, NGCircos, CircosJS, JBrowse 2 and Cosmograph |
| Programmatic access | *Not applicable* | *Not applicable* | No | API and Cytoscape app | Documented REST API, Swagger UI, code in four languages, worked examples tested automatically |
| Undo and redo | No | No | No | No | Every scene change |
| Rendering control | No | No | No | Frame rate | Frame rate, resolution, node detail, presets and live statistics |
| Privacy | *Not described* | *Not described* | *Not described* | *Not described* | No login or cookies, data processed but not stored, requested links deleted after one hour |
| Security hardening | *Not described* | *Not described* | *Not described* | *Not described* | Input validation, size and time limits, content security policy, dependency and code scanning |
| Documentation and examples | Help pages | Help pages | Help pages | Help pages, example sessions | A help page per panel, recipes, API and privacy pages, examples including six neural networks |

## MATERIALS AND METHODS

### Implementation and architecture

Arena3D 3.0 replaces the R and Shiny code of Arena3D^web^ with a stateless Python service and a browser client. The FastAPI service reads networks and sessions and computes layouts, communities, metrics, plots, perturbations, enrichment and database imports. Graph algorithms come from the Python interface of igraph (21), which shares its C core with the R package used previously. Multiplex communities are computed with the leidenalg implementation of the Leiden algorithm (40), and UMAP with its reference implementation (41). On the public server, a network can hold up to 20 layers, 10 edge channels, 20,000 nodes, each counted once in every layer it belongs to, and 10,000 edges. Kamada-Kawai, multidimensional scaling, Graphopt and Sugiyama accept up to 2,000 nodes and UMAP up to 5,000. Supplementary Table S2 gives the run times of the layouts, metrics, community detection, attacks and edge bundling on networks of 1,000 to 10,000 edges. The client is written in TypeScript and uses three.js for rendering, with three rendering presets and live rendering statistics. Every change to the scene can be undone and redone. On phones and tablets the mouse is replaced by touch gestures. A help page for each panel and step-by-step recipes for common tasks are built in.

### Input and database import

Networks are loaded in the Arena3D tab-separated format, which lists the source node, source layer, target node and target layer of each edge together with an optional weight and channel. Two-column edge lists and GraphML (42), GML, SIF and CX2 files with a layer attribute are also accepted. Separate attribute files can add colors, shapes, labels, descriptions and annotation terms, NORMA group files add groups, and numeric tables add value columns such as expression levels.

Six importers fetch networks from public databases (43, 44). For STRING (45), the organism is chosen from a list of 19 or given as an NCBI taxon identifier, and functional or physical networks can be retrieved with up to 500 added interactors and a chosen minimum confidence. Interactions can be split into evidence channels, enriched terms from 21 collections are turned into annotation sets, and a versioned STRING address keeps the results reproducible. IntAct (46) and OmniPath (47, 48) interactions are retrieved for the same proteins, either alone or as extra layers next to STRING with the copies of each protein linked. NDEx networks (49) are loaded by identifiers, while Reactome pathways (50) and Gene Ontology terms (51) become layers of their genes, connected by the STRING interactions among them. Before each load, the user chooses Multilayer or Single network mode, whether the network replaces or extends the current one, and whether it is laid out at once with the Fruchterman-Reingold algorithm (52).

### Layouts in two and three dimensions and alignment

Twelve layout algorithms are available. Eleven of them are the igraph layouts used in Arena3D^web^, including Fruchterman-Reingold, Kamada-Kawai (53), DrL (54), Reingold-Tilford (55) and Sugiyama (56), and the twelfth is UMAP (57). A layout can be applied to each selected layer separately, to the selected layers as one network, or only to selected nodes, and edge and channel weights can be considered. Seven of the algorithms (Fruchterman-Reingold, Kamada-Kawai, DrL, random, grid, circle on a sphere and UMAP) also work in 3D, which lifts the nodes off their floor or fills a container. UMAP positions nodes by topology, by chosen attributes such as value columns, annotation terms, clusters and metrics, or by a weighted combination of the two, and a fixed seed makes the result reproducible. For topology, each node is represented by its probabilities of reaching the other nodes in one or two steps of a random walk. In Single network mode the layers are merged into one graph and laid out in 3D without floors, and a Spread slider opens up dense regions by scaling distances from the center. Exact alignment merges the selected layers by node name, lays them out once and places each copy of a node at the same position on every floor, so the links between copies run straight. Three pseudo-alignment modes balance alignment against the structure of each layer with a strength between 0 and 1. Soft alignment turns the layout of each layer, mirrored if that fits better, to lie as close as possible to the exact layout (58) and then blends the two, from the turned layout alone at 0 to the exact layout at 1. Anchor alignment starts each layer from the positions of its nodes in a reference layer and pulls them toward these positions in proportion to the strength, pinning them at 1. Chain alignment does the same with each layer anchored to the one before it. Each multilayer layout reports a misalignment score, the mean distance within the layer planes between linked copies as a percentage of the layout diagonal. In the alignment benchmark (Results), each mode was run with Fruchterman-Reingold and ten random seeds, and the structure kept in each layer was measured as the Spearman correlation ρ between drawn and shortest-path distances, averaged over the layers. Independent layouts keep each layer at its own size in the app, so in the benchmark they were scaled to the frame that the aligned modes share.

### Layer geometry and 3D containers

A layer can be shown as a square or round floor, with no floor at all, or as one of thirteen 3D containers. Eleven of these are geometric solids (cube, sphere, cylinder, cone, pyramid, dome, ellipsoid, torus, hexagonal prism, geodesic sphere and disc stack), and the other two are softer shapes, a fog cloud modeled on the group clouds of NORMA and a bubble. Nodes in a container receive a 3D layout within its volume or on its surface. In cones, pyramids and disc stacks the nodes are placed in tiers by degree, and the torus wraps a floor layout around a ring. Color, opacity, outline, wireframe and glow can be set for each layer, floors come in five styles, and each layer can display its own coordinate axes.

The layers themselves can be arranged in thirteen ways, ten of them geometric (parallel planes, zigzag, star, cube, helix, sphere, grid, circle, stack and fan). A force layout can group heavily linked layers, a chord arrangement places the floors on a ring with the edges between layers as chords, and the layers can also be drawn as a hive plot (38). In a hive plot, each layer becomes an axis radiating from a common center, and each node is placed on its axis by degree, betweenness, clustering coefficient or name. Arena3D 3.0 can also spread the axes through space, as a tripod for three layers, a tetrahedron for four, the six coordinate directions for six and evenly over a sphere for other numbers.

### Clustering, group annotations and group layouts

Communities can be detected with Louvain (59), Walktrap (60), fast greedy modularity optimization (61), label propagation (62) or Leiden (40). They are computed within each layer, on the selected layers merged into one graph, or across layers by node name with multiplex Leiden, and the result can be stored as an annotation set. Annotation sets can also be loaded from files or from STRING, and one network can carry several of them. In the Annotations panel of the View menu, each group is drawn around its members in one of seven styles, as a hull or outline on the floor, a fog cloud, a cloud of puffs, or a translucent sphere, ellipsoid or box. Nodes that belong to several groups are drawn as pie charts. Groups can be hidden individually, and their statistics and overlaps are reported.

The three group layouts of NORMA 2.0 (37) were reimplemented with their original weighting schemes. With a *virtual node* per group, a hidden hub is linked to all members of the group. *Group gravity* strengthens the edges inside groups and weakens the edges between them. *Super nodes* first collapse each group into a single node, lay out the collapsed graph and then lay out each group within its region. The groups can be clusters, annotation terms or the layers themselves. In the last case the whole network is laid out in 3D, and each floor moves to the center of its group and turns away from the middle of the arrangement.

### Edge bundling

Force-directed edge bundling (63) was implemented in 3D. Each edge is divided into a chain of points connected by springs, and points on compatible edges, those with similar direction, length and position, attract each other until related edges merge into common bundles. Edges within and between layers are bundled with separate strengths, and in Single network mode all edges are bundled together. Because the bundles do not depend on the camera, the scene can be rotated without recomputing them, and the channels of an edge run side by side along their bundle. Bundling runs in the browser and took 2.3 s for 10,000 edges on a laptop with an Apple M3 Pro processor (Supplementary Table S2).

### Analysis including multilayer metrics, perturbation and plots

Nodes can be sized and colored by degree, clustering coefficient, betweenness, closeness, eigenvector centrality, PageRank, coreness or eccentricity. These metrics are computed per layer or on the selected layers, for undirected, directed or multiedge graphs, as in the Network Analysis Profiler (64, 65). Multiplex degree, the participation coefficient (66), which is high for nodes whose links are spread evenly over the layers, and multiplex PageRank (67) are provided as multilayer measures. Network statistics include density, degree, components, diameter, path length, clustering, transitivity and assortativity for each layer and for the whole network. A layer similarity view reports the Jaccard overlap of nodes and edges between layers, and selected nodes can be submitted to STRING for enrichment analysis.

The Plots panel shows the distribution of any metric, edge weight, value column, component size or path length. Distributions are drawn as histograms, density curves or cumulative distributions, or on logarithmic axes with an optional power-law fit (68), and box and violin plots compare the layers. Other plots relate topological properties, for example scatter plots with Pearson and Spearman correlations, a correlation matrix of all metrics, the clustering coefficient and the mean neighbor degree as functions of degree, the rich-club coefficient normalized against degree-preserving randomized graphs (69), k-core shells, centrality rankings across layers and layer profiles. Layers are overlaid in their own colors, charts can be saved as SVG, PNG or CSV, and clicking a bar or point selects the corresponding nodes.

In the Perturbation panel, nodes can be removed by selection, by name or as the top-ranked nodes by degree, betweenness or PageRank, optionally together with their copies in other layers. Edges can be removed by selection, by channel, between layers or at random. Each layer and the whole network are then compared before and after the knockout in terms of nodes, edges, components, density, path length, diameter, global efficiency (70) and isolated nodes. A table lists the change in degree, betweenness and harmonic closeness of every node, which can also color the nodes. Network propagation by random walk with restart (71) can be compared before and after as well. In attack mode, nodes are removed at random or in order of degree or betweenness, ranked once or recalculated after each step, and the relative size of the largest component is plotted for each layer and for the whole network together with the robustness measure R (72).

Nodes are selected by double click, with a rectangle or a freehand lasso, or by component, layer or degree. In a live mode, the shortest paths among the selected nodes are highlighted within and across layers. The Find panel searches nodes and edges by exact, prefix, suffix, substring or regular expression match, lists the hits in tables and can focus the camera on them.

### Styling, editing and labels

Nodes can take one of fourteen shapes, and the channels of an edge are drawn as parallel lines, as on the STRING website. Arrowheads can be limited to the directed layers, so that undirected interactions such as those of STRING can be shown next to directed signaling. Nodes and edges are styled through property tables modeled on the Style panel of Cytoscape (16), applied to the selected nodes, all nodes or a single layer. Node properties include color, opacity, border, size, proportions, shape, label and description, and edge properties include color, width, opacity, line type and arrow color. An attribute editor changes a single cell or all checked rows at once, stays in sync with the selection in the scene and exports attribute files that load back, and an inspector lists every attribute of a node or an edge. By default, labels are drawn with a thin contrasting halo, and box, shadow, plain and bold styles and four label positions are also available. Label colors can follow layers, clusters or annotation terms, and a label picker sets which labels stay visible.

Nodes are colored by layer, cluster, annotation, metric or value column, with categorical palettes or sequential palettes such as cividis (73). A theme sets the background, floors, nodes, edges, channels, labels and containers at once, and the 27 themes are grouped into dark, light and color-blind safe themes. The last group contains the Okabe-Ito palette (74), three palettes by Paul Tol (75), viridis (76) and three ColorBrewer sets (Set2, Dark2 and Paired), which ColorBrewer (77) marks as color-blind safe for up to three or four categories. In every theme, each layer and each of the ten channels receives a color with a contrast ratio of at least 3 against the background, the minimum that the Web Content Accessibility Guidelines (78) set for graphics. A color-blind mode also gives each layer its own node shape, which keeps the layers apart when there are more of them than safe colors, and a preview simulates protanopia, deuteranopia and tritanopia (79), and achromatopsia as luminance alone. When the theme is changed while a network is open, the user can choose whether the network keeps its own colors.

### Alternative views and exports

A chord diagram in the style of Circos (80) shows the layers as arcs and the edges as chords. A Sankey diagram (81) shows the layers as columns and the edges between layers as flows, banded by node, group or layer, and an adjacency heatmap groups the nodes by layer. Layer comparison, group overlap and the comparison of two networks are shown as interactive UpSet plots (39). Bars or sets can be picked as a union or an intersection and then selected in the scene or loaded as a new network. A UMAP view shows the nodes as a scatter plot and exports their coordinates as CSV, and all reports are given as sortable tables.

The view can be exported as PNG, SVG, a GLB 3D model or a rotation video, and the network as GraphML, CX2, Cytoscape.js JSON (20), SIF, MuxViz files (27) or a JSON session. Four browser-based alternatives to Circos are also provided. These are a Gosling (82) specification, pages for NGCircos (83) and CircosJS (github.com/nicgirault/circosJS) with degree or value tracks, and a JBrowse 2 (84) configuration in which layers act as sequences and edges as breakends drawn as chords. Node and edge tables can also be sent to Cosmograph (cosmograph.app).

### Integration and the REST API

A network can be opened in a running Cytoscape session through cyREST (85), in Gephi Lite (17) through a GraphML file hosted for one hour, in the Vega editor (86) as an interactive heatmap, on the STRING website or in NORMA. The genes of selected layers can be sent to Flame (87, 88) for enrichment analysis. In the other direction, applications can send sessions to Arena3D through the REST API and receive a link that opens them, which is how networks are passed from Cytoscape to Arena3D^web^ (35). A request to /api/open converts a plain edge list with optional node colors, sizes and annotations into such a link and lays the network out. All JSON endpoints are documented in an OpenAPI description, with an example for each endpoint that the automated tests check. An API menu adds example code in curl, Python, R and JavaScript and an embedded Swagger UI for trying every endpoint.

### Privacy and availability

Arena3D 3.0 is free and open to all users, without login or registration. Networks are processed on the server but not stored. Only links that a user requests, such as hosted exports and API links, are kept under random tokens and deleted after one hour. No cookies, analytics or tracking are used, and external services such as STRING are contacted only on request. Uploads and computations are validated and limited in size and time, a content security policy restricts what the pages may load, and the code and its dependencies are scanned for known vulnerabilities. The application runs in current versions of Chrome, Edge, Firefox and Safari, and its interface is tested automatically in Chrome on a desktop screen and in emulations of a phone and a tablet.

### Datasets

Two real networks are distributed with the application. The first is the SARS-CoV-2 and host protein interaction network of Gordon *et al.* (89), with 455 nodes and 985 edges in five layers of viral proteins, host proteins, drugs, complexes and biological processes. The second is the human GPCR signal transduction network (90), with 63 nodes and 332 edges in four layers of GPCRs, G proteins, GRKs and effectors. Both were used before with Arena3D^web^ (34, 35). Additional real networks were retrieved for this article from STRING version 12.5 (45), whose enrichment analysis provided the KEGG pathways (91) used as groups, from IntAct (46) and from OmniPath (47, 48). The STRING networks of Figure 3 are human, the functional one of Figure 3F with a minimum score of 0.4 and the physical one of Figure 3G with a minimum score of 0.7. The alignment benchmark uses two human networks of three layers, built from STRING interactions with a combined score of at least 0.4, IntAct interactions with an MI score of at least 0.4 and all OmniPath interactions. The first links 94 proteins of the cell cycle, the DNA damage response, apoptosis and growth factor signaling, and the second 81 proteins of innate immune signaling. Both protein lists were selected for this article (Supplementary Table S1, with the KEGG pathways of each protein). The first network has 94, 91 and 94 proteins with 2,590, 221 and 524 interactions in its STRING, IntAct and OmniPath layers and 185 links between copies, and the second 81, 70 and 76 proteins with 2,261, 140 and 235 interactions and 146 links. The first network was also used for the robustness and degree analyses (Results).

The Six layers example from the Arena3D^web^ article (34) contains six layers with random topologies of 94 to 558 nodes and 91 to 740 edges each, with 1,600 nodes and 2,396 edges in total. The synthetic multiplex networks were generated for this article. They contain 90 genes in six pathway modules of 15 genes each (cell cycle, MAPK, PI3K-AKT, DNA repair, apoptosis and Wnt signaling), each gene in one module, on four layers (co-expression, protein interactions, genetic interactions and regulation) or six (adding phosphorylation and pathway membership). In these layers each gene is present with a probability of 0.92, 0.88, 0.80, 0.85,

0.70 and 0.95, two genes of the same module are linked with a probability of 0.42, 0.34, 0.26, 0.20, 0.30 and 0.38 and genes of different modules with a probability of 0.003, 0.004, 0.005, 0.002, 0.003 and 0.001. Three to six hub genes in each of the protein interaction, genetic interaction, regulation and phosphorylation layers are linked with 1.8 times these probabilities, plus 0.03 across modules, and about 40% of the genes are linked to their copies in the next layer. To test community detection on weaker modules, 20 four-layer networks were also generated for each mixing μ from 0.05 to 0.8, the expected share of a gene’s links that leave its module, at the same expected degree per layer. Their communities were compared with the planted modules by the adjusted Rand index (92). The two small networks used for the comparison are the protein interactions among the cell cycle and DNA repair genes (network A, 28 genes and 68 edges) and the genetic interactions among the apoptosis and DNA repair genes (network B, 22 genes and 38 edges) of the four-layer network. The gene symbols are real, but the links are random, so the rankings and knockouts computed on these networks illustrate the tools and carry no biological meaning. The six neural network examples are modeled on LeNet (93), the transformer (94), the long short-term memory network (95), U-Net (96), the variational autoencoder (97), and a graph convolutional network (98) applied to the karate club network of Zachary (99). They illustrate architectures and are not trained models.

## RESULTS

### The visual repertoire

Layers do not have to share a layout or a shape (Figure 1A to 1E), and modules can be drawn as fog clouds or laid out apart on floors and inside containers (Figure 1F to 1J). Figure 1K to 1M show edge bundling, a helix arrangement and Single network mode.

**Figure 1.**
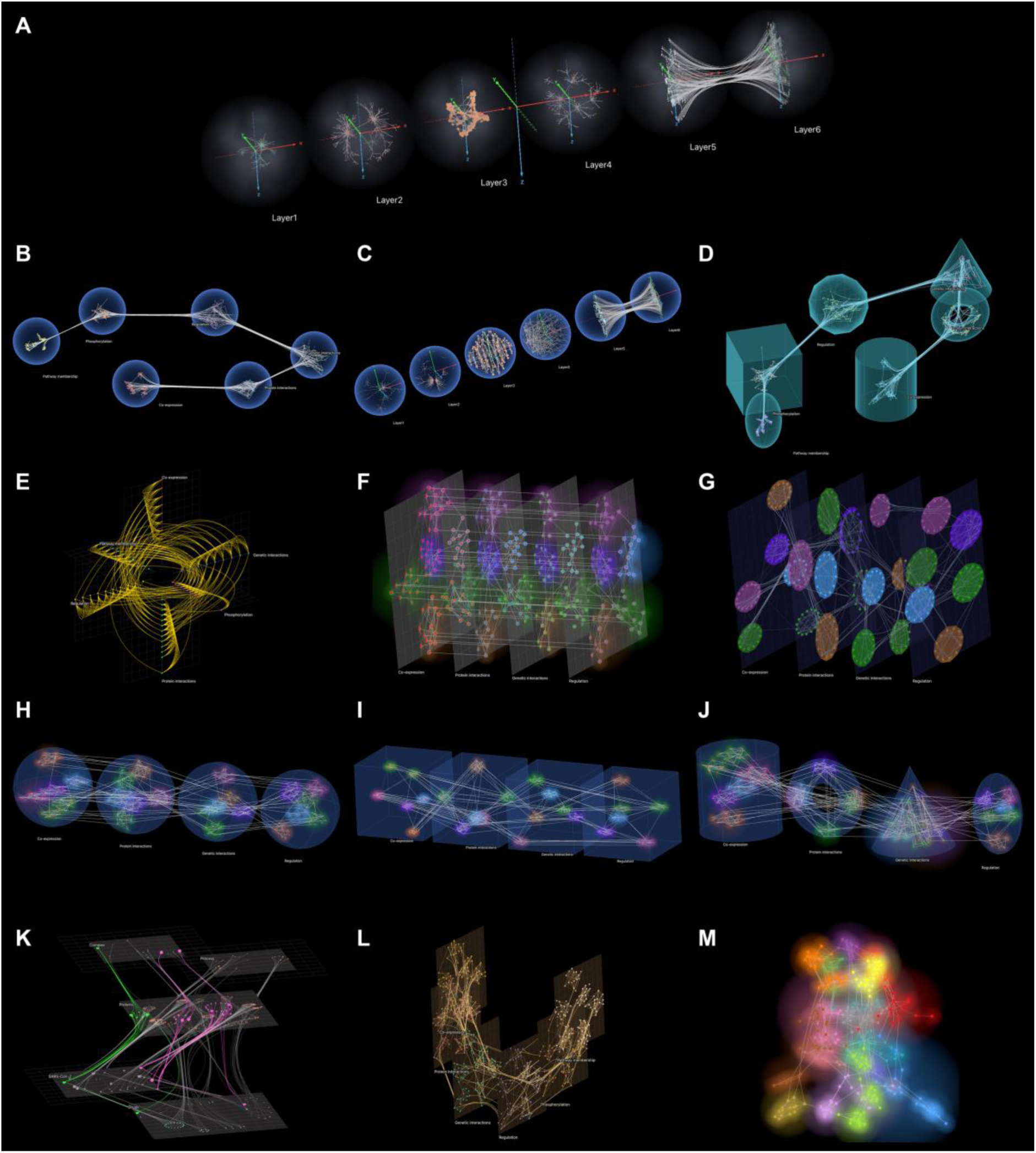
The visual repertoire of Arena3D 3.0. All panels are drawn on a black background. (**A**) The Six layers example (six random topologies, 1,600 nodes) in glowing spheres with their own axes, the links between ‘Layer5’ and ‘Layer6’ bundled. (**B**) The six-layer synthetic network with its spheres on a ring. (**C**) The Six layers example with Fruchterman-Reingold, Kamada-Kawai, grid and circle on a sphere layouts on its first four layers and its last two layers aligned. (**D**) Cylinder, torus, cone, geodesic sphere, cube and ellipsoid containers in the Ripple theme. (**E**) A hive plot of the six-layer synthetic network, the nodes placed by degree, in the High contrast theme. (**F**) The four-layer synthetic network, its modules as fog clouds and each node ringed in the color of its module. (**G**) A virtual node group layout with the groups outlined by hulls, in the Universe theme. (**H** to **J**) Virtual *nodes* in spheres (**H**), *group gravity* in cubes (**I**) and *super nodes* in a cylinder, a torus, a cone and an ellipsoid (**J**). (**K**) The SARS-CoV-2 network with its edges bundled within and between layers. (**L**) The network of panel B as a helix in the Marquee theme. (**M**) The network of panel B in Single network mode, its Louvain communities as fog clouds.

### Analysis, editing and interaction

Figure 2 shows the analysis and editing tools on the synthetic networks and network A, with annotation groups and multiplex communities (Figure 2A and 2B), the participation coefficient (Figure 2C), a knockout of the two genes of highest betweenness in network A (Figure 2D), centrality rankings and layer profiles (Figure 2G and 2H), the attribute table (Figure 2I) and live shortest paths (Figure 2J). Over 20 random seeds, multiplex Leiden recovered the six planted modules on all 90 genes, with an adjusted Rand index of 1.0. Leiden on single layers, on the genes each layer contains, scored 1.0 on co-expression and protein interactions and on average 0.91 and 0.84 on the sparser genetic interaction and regulation layers. With weaker modules (Datasets), multiplex Leiden still scored 0.99 and 0.96 when 30% and 40% of the links left their module, while single layers fell to 0.38 to 0.86 and 0.24 to 0.62. At a mixing of 0.6 or more no method recovered the modules (Figure 2K).

**Figure 2.**
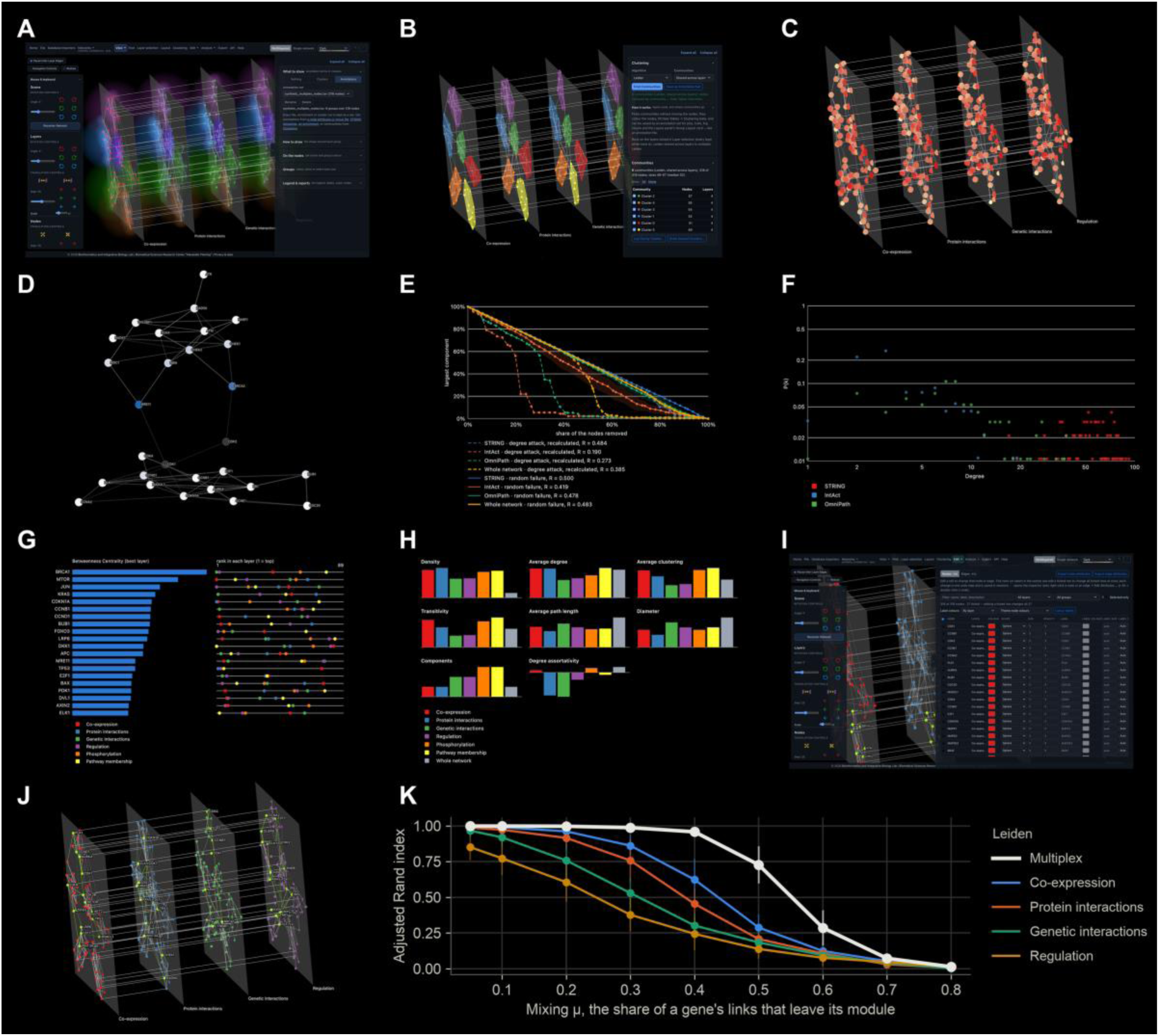
Analysis, editing and interaction. Panels A to C, I and J show the synthetic network of four layers, D network A of Figure 3J, E and F the DNA damage network of the alignment benchmark, G and H the synthetic network of six layers and K synthetic networks with weaker modules. (**A**) The interface with the Annotations panel of the View menu next to the scene, the annotation groups drawn as fog clouds. (**B**) Communities found with multiplex Leiden, shared across layers and drawn as hulls, next to the summary of the Clustering panel. (**C**) Nodes scaled and colored by their participation coefficient, from low (pale) to high (red). (**D**) A knockout of CDK1 and CDK2, the two genes of highest betweenness in network A, shown faded, with the other genes colored by their change in betweenness (blue lost, red gained, pale unchanged). (**E**) Robustness curves of the STRING, IntAct and OmniPath layers and of the whole network under degree-targeted attack with recalculated ranks (dashed lines) and random failure (solid lines), with the robustness measure R of each curve. (**F**) Degree distributions of the three layers on logarithmic axes. (**G**) Centrality ranking across six layers, the nodes of highest betweenness in their best layer with their rank in every layer. (**H**) Layer profiles of six layers and the whole network, with density, average degree, average clustering, transitivity, average path length, diameter, components and degree assortativity. (**I**) The attribute table of the Edit menu, in which one or many rows are edited at once, with the selected DNA repair genes highlighted and labeled in the scene. (**J**) Live shortest paths among CDK1, TP53, CTNNB1 and KRAS within and across layers, with the nodes along them and halo labels. (**K**) Adjusted Rand index of multiplex Leiden and of Leiden on each layer against the planted modules of four-layer synthetic networks, as the share of links between modules (the mixing μ) grows, means of 20 networks with one standard deviation.

On the DNA damage network of the alignment benchmark (Datasets), degree-targeted attack, with the ranks recalculated after each removal, lowered R to 0.19 for IntAct, 0.27 for OmniPath and 0.39 for the whole network, against 0.42, 0.48 and 0.48 under random failure (Figure 2E). The dense STRING layer kept 0.48 against 0.50. The degree distributions of this network are drawn without a power-law fit (Figure 2F), because the method of Clauset et al. (68) found no layer whose tail a power law fitted better than an exponential or a lognormal distribution.

### Alternative views, exports and integration

Figure 3 shows the chord, Sankey, heatmap, UpSet and UMAP views of the synthetic networks (Figure 3A to 3E), multi-edged STRING and directed OmniPath layers from the importers (Figure 3F), enriched pathways as annotation groups (Figure 3G), an export to NORMA (Figure 3H), the interface on a phone and a tablet (Figure 3I) and the comparison of networks A and B (Figure 3J). Among the twelve proteins of Figure 3G, 47 KEGG pathways were enriched against the human genome at a false discovery rate below 0.05. Homologous recombination and p53 signaling, the DNA repair and damage response pathways among the three most enriched, were drawn as groups. ATM, which STRING assigns to both, is drawn as a pie chart between them.

**Figure 3.**
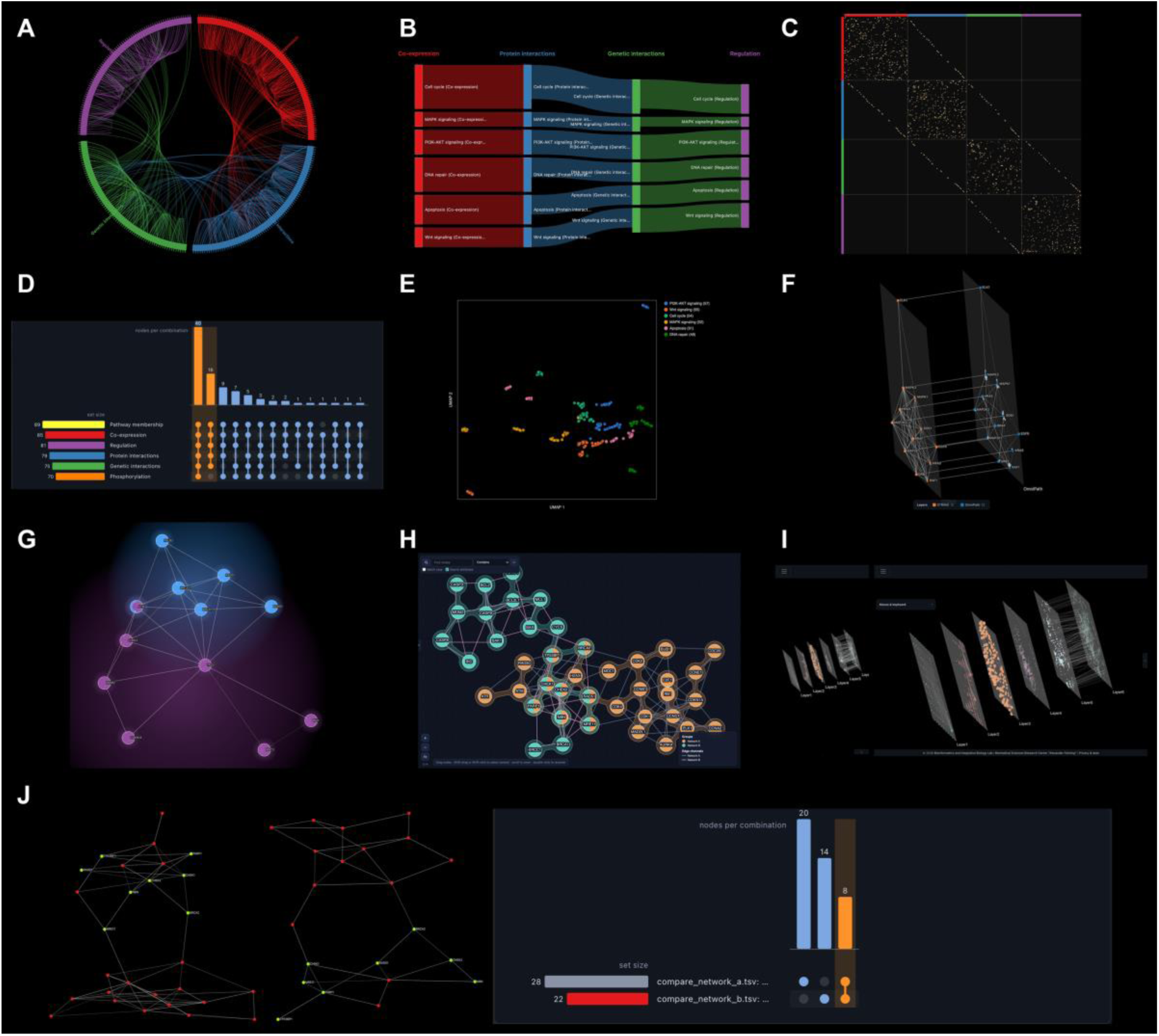
Alternative views, exports, devices and integration. (**A**) Chord diagram of the network of four layers, with the layers as arcs and the edges as chords. (**B**) Sankey diagram of the edges between layers, banded by pathway module. (**C**) Adjacency heatmap ordered by layer, with the edges within layers in the diagonal blocks and the links between the copies of a node on lines parallel to the diagonal. (**D**) Interactive UpSet plot of the node overlap between six layers, with two selected intersections highlighted. (**E**) UMAP view of the 318 layer copies of the 90 genes by topology, colored by pathway module. (**F**) A multi-edged and a directed network from the database importers, STRING interactions among twelve proteins of EGFR and MAPK signaling (EGFR, GRB2, SOS1, HRAS, KRAS, RAF1, BRAF, MAP2K1, MAP2K2, MAPK1, MAPK3 and ELK1) with one edge per evidence channel, and directed OmniPath signaling among the same proteins added as a second layer with arrowheads on its edges only, the copies of each protein linked across the layers. (**G**) A STRING network of twelve proteins (BRCA1, BRCA2, RAD51, PALB2, BARD1, ATM, CHEK2, TP53, MDM2, CDKN1A, BAX and BBC3) with two of its 47 enriched KEGG pathways, homologous recombination and p53 signaling, drawn as fog clouds and ATM, which STRING assigns to both, as a pie chart between them. (**H**) Networks A and B loaded as two layers and exported to NORMA, where each layer becomes an annotation group and the eight genes found in both are drawn as pie-chart nodes between the groups. (**I**) The Six layers example on a phone and on a tablet. (**J**) Networks A and B compared, each with the eight genes they share selected and labeled, next to the UpSet plot of their node overlap.

### Themes and color vision

Figure 4 shows sixteen of the 27 themes, including the Universe, Marquee and Ripple styles of Cytoscape and the eight palettes of the color-blind group.

**Figure 4.**
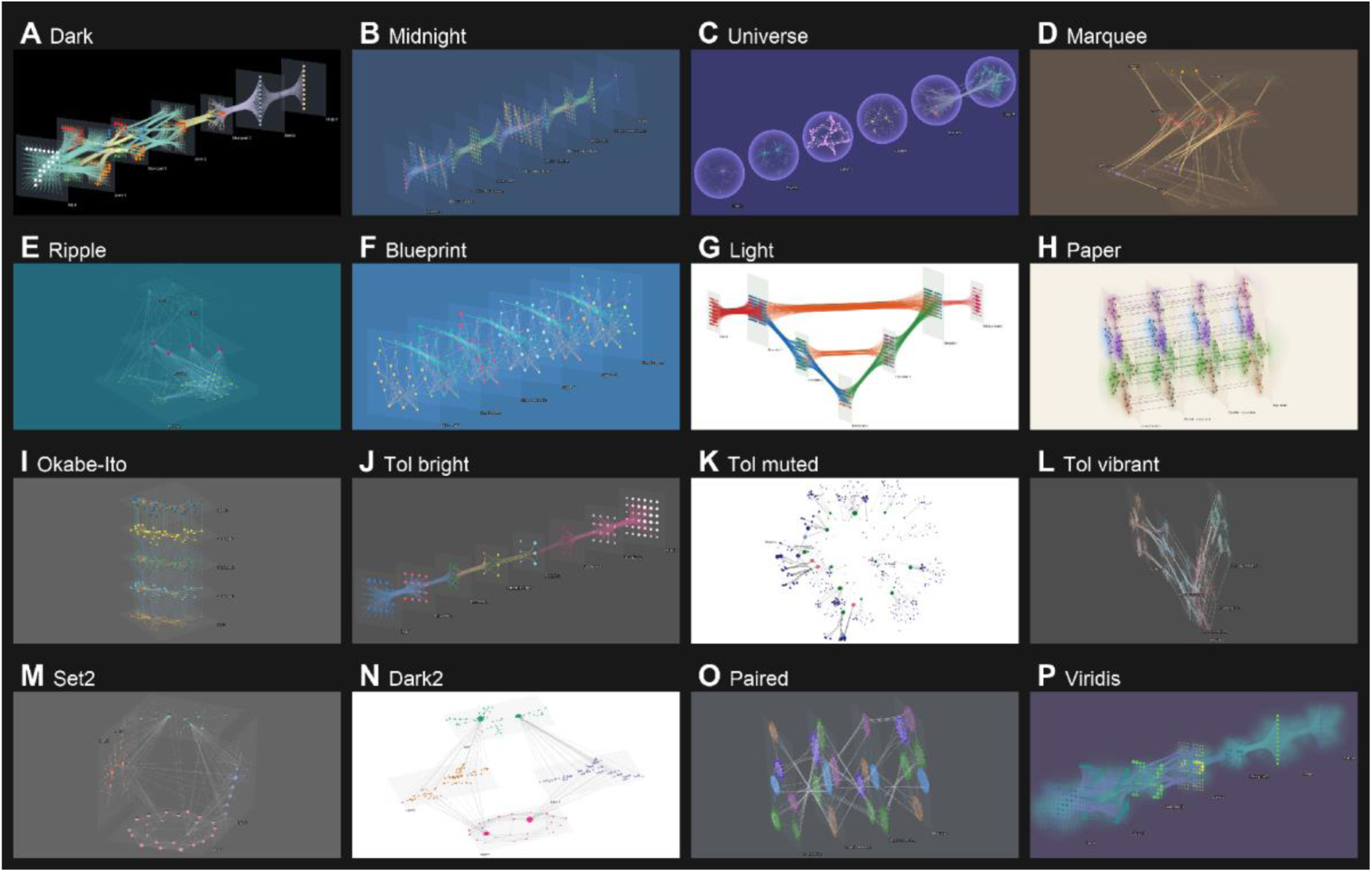
Themes and color vision. Sixteen of the 27 themes, each applied with its own background and colors to a different example network. (**A**) Dark with the colors of the example itself, the LeNet-type convolutional network reading a drawn digit 7 in its input layer. (**B**) Midnight, the transformer. (**C**) Universe, the Six layers example in glowing spheres. (**D**) Marquee, the SARS-CoV-2 network. (**E**) Ripple, the GPCR signal transduction network of four layers. (**F**) Blueprint, a long short-term memory network unrolled over six time steps. (**G**) Light, a U-shaped segmentation network. (**H**) Paper, the pathway modules of the synthetic network of four layers as fog clouds. (**I** to **P**) The eight color-blind safe palettes, Okabe-Ito on a graph neural network over the karate club network, Tol bright on a variational autoencoder, Tol muted on the SARS-CoV-2 network in Single network mode, Tol vibrant on the synthetic network of six layers as a helix, Set2 on four layers forming a cube, Dark2 on four layers with node and edge attributes, Paired on the synthetic network of four layers with a group layout, its modules outlined by hulls, and viridis on the convolutional network with its layers as clouds.

### Alignment of node copies across layers

Alignment was benchmarked on the two STRING, IntAct and OmniPath networks described in Datasets (Figure 5). Over ten seeds, independent layouts left the copies of a protein on average 26% and 29% of the layout diagonal apart, with ρ of 0.68 and 0.72 (Figure 5D and 5E). Figure 5A to 5C show one seed. Exact alignment removed this distance and lowered ρ to 0.47 and 0.32 (Figure 5C). Soft alignment at strength 0.5 reduced misalignment to 10% and 12% with ρ of 0.59 and 0.54 (Figure 5B). At strength 0, where each layer is only turned onto the exact layout, misalignment fell to 17% and 21% without lowering ρ. Anchor and chain alignment kept most of the structure only at strength 0.25, where misalignment stayed at 21% to 26%. At 0.75 they left the copies 8% to 12% of the diagonal apart while keeping no more structure than exact alignment, with ρ of 0.40 and 0.43 against 0.47 and of 0.31 and 0.32 against 0.32.

**Figure 5.**
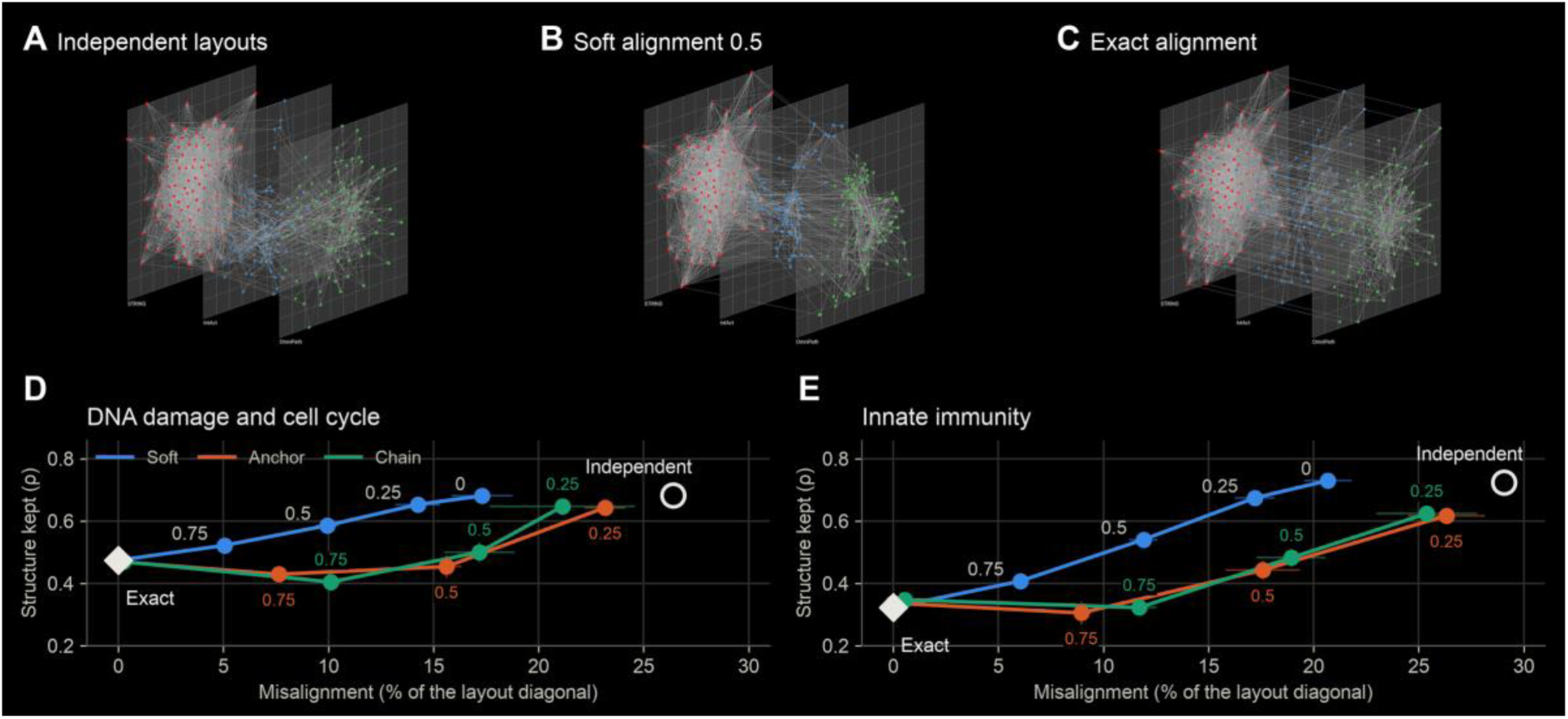
Alignment of node copies across layers. (A to C) The DNA damage and cell cycle network (94 proteins in STRING, IntAct and OmniPath layers) for the default seed, with independent Fruchterman-Reingold layouts of the three layers (A), soft alignment at strength 0.5 (B) and exact alignment (C), with a misalignment of 23.8%, 10.8% and 0% and ρ of 0.69, 0.57 and 0.48. (D and E) ρ, the Spearman correlation between drawn and shortest-path distances averaged over the layers, against misalignment for soft, anchor and chain alignment at the strengths labeled next to the points, on this network (D) and on the innate immunity network of 81 proteins (E). Points are means of ten seeds with one standard deviation, the open circle marks independent layouts drawn at the same size and the diamond exact alignment.

## DISCUSSION

Table 1 compares Arena3D 3.0 with the four earlier releases. The 2008 and 2012 versions were Java desktop applications that are no longer distributed, and the two Arena3D^web^ releases ran on an R and Shiny server with up to 10 layers, all on flat floors. Arena3D 3.0 holds up to 20 layers on the public server and adds database import, perturbation analysis and undo, which no earlier release offered.

Among general tools, Cytoscape (16) and Gephi (17) offer many more plugins and layouts for single networks. Arena3D 3.0 follows the style conventions of Cytoscape and can pass networks to both tools, but neither uses the layers of a network as geometry. MuxViz (27) provides more multilayer measures in a locally installed application, and Arena3D 3.0 exports to its format.

Although it was designed with biological data in mind, Arena3D 3.0 can be used for any system of typed entities and relations. In the six neural network examples, for instance, the layers of a network map onto Arena3D layers and its connection types onto channels (Figure 4).

### Limitations

Networks above the limits of the public server require a local installation, where the limits can be raised. Pseudo-alignment and group layouts work in two dimensions within each layer, except inside 3D containers and when the layers themselves are the groups. The chord and Sankey diagrams are static drawings, and the heatmap becomes interactive only in the Vega editor. The virtual reality mode of the second release has not yet been ported to the new client.

## DATA AVAILABILITY

Arena3D 3.0 is freely available at www.arena3d.org and runs in current web browsers without plugins, login or registration. The source code is available at github.com/Pavlopoulos-Lab/Arena3D and a Docker image (pavlopouloslab/arena3dweb on Docker Hub, which keeps the name of the previous release) allows local deployment with adjustable limits. The REST API is documented at www.arena3d.org/api/docs. The SARS-CoV-2, GPCR, Six layers and neural network examples are distributed with the application and listed with their files on its Examples help page. The synthetic and benchmark networks, the scripts that generated and analyzed them and Supplementary Tables S1 and S2 are provided as Supplementary Data.

## SUPPLEMENTARY DATA

Supplementary Data are available at NAR Online.

## ACKNOWLEDGEMENTS

Claude Opus 5.5 (Anthropic) assisted with optimizing the source code and with drafting and editing the manuscript. The authors reviewed and verified all content and take full responsibility for it.

