## Supplementary material for "Arena3D 3.0: a web server for exploring, analyzing, annotating and sharing multilayered networks in 3D": README.txt

Arena3D 3.0, Supplementary Data

Networks
  synthetic_multiplex.tsv, synthetic_multiplex_nodes.tsv  the four-layer synthetic network (mock_data.py, seed 7)
  mock6_multiplex.tsv, mock6_multiplex_nodes.tsv          the six-layer synthetic network (mock_data6.py)
  compare_network_a.tsv, compare_network_b.tsv            networks A and B (Figures 2D, 3H and 3J)
  alignment_network.json/.tsv, alignment_network_immune.json
                                                          the two benchmark networks (STRING, IntAct, OmniPath)

Scripts (Python 3.12 or later; the analyses import the Arena3D backend,
github.com/Pavlopoulos-Lab/Arena3D, from ../backend)
  mock_data.py, mock_data6.py   generate the synthetic networks
  alignment_benchmark.py        the alignment benchmark of Figure 5 (needs the backend on localhost:8000)
  alignment_panel.py            the charts of Figure 5D and 5E
  rob4_analyses.py              ARI under weaker modules, robustness and degrees of the DNA damage network,
                                the enrichment of Figure 3G, Table S1 and the run times of Table S2
  degree_fit.py                 the power-law test of the degree distributions (Clauset et al. 2009)
  fdeb_time.mts                 the run time of the client's edge bundling (node --experimental-strip-types)
  ari_panel.py                  the chart of Figure 2K

Results
  alignment_benchmark*.csv, rob4_*.csv, rob4_degrees.json, Table_S1_benchmark_proteins.tsv
