## Supplementary material for "Arena3D 3.0: a web server for exploring, analyzing, annotating and sharing multilayered networks in 3D": alignment_benchmark.py

```
"""Alignment benchmark for the NAR manuscript (Rob review, round 3, analysis 1).

A real multilayer network, STRING, IntAct and OmniPath interactions among the
same human proteins as three layers with the copies of each protein linked
(POST /api/import/databases), is laid out with Fruchterman-Reingold under every
alignment mode of Arena3D 3.0: each layer on its own, soft, anchor and chain
alignment at several strengths, and exact (coupled) alignment. For every run
the app's own misalignment score is recorded (POST /api/layout), together with
how well each layer keeps its own structure: the Spearman correlation between
distances in the drawing and shortest-path distances in the layer's graph,
averaged over the layers. Ten seeds per mode.

The plain per-layer layout keeps each layer's own size, as the app draws it, so
the baseline without alignment is that layout with every layer scaled to the
common frame the alignment modes use ("Independent, same size"). Soft
alignment turns each layer onto the aligned layout before blending (since the
fix of 2026-10-23); the results from before the fix are kept in
data/alignment_benchmark*_before_fix.csv.

Needs the backend on localhost:8000. The imported network is saved once to
data/alignment_network*.json and reused, so the numbers can be repeated.

Run: ../backend/.venv/bin/python alignment_benchmark.py
"""

import csv
import json
import statistics
import sys
import urllib.request
from itertools import combinations
from pathlib import Path

import igraph as ig
from scipy.stats import spearmanr

sys.path.insert(0, str(Path(__file__).parent.parent / "backend"))
from app.models.network import EdgeModel, NodeModel  # noqa: E402
from app.services.alignment import _fit, misalignment  # noqa: E402

API = "http://localhost:8000"
HERE = Path(__file__).parent
DATA = HERE / "data"
SEEDS = range(1, 11)

# two gene sets: cell cycle, DNA damage response, p53 and apoptosis and growth
# factor signaling (the main one), and innate immune and cytokine signaling (a
# second network, to see whether the result holds)
GENES = """
CDK1 CDK2 CDK4 CDK6 CCNA2 CCNB1 CCND1 CCNE1 CDC20 CDC25A CDC25C CDKN1A CDKN1B CDKN2A
RB1 E2F1 PLK1 AURKA AURKB BUB1 BUB1B MAD2L1 CHEK1 CHEK2 WEE1 SKP2
ATM ATR BRCA1 BRCA2 RAD51 PALB2 BARD1 MRE11 RAD50 NBN H2AX TP53BP1 MDC1 PARP1 XRCC1
XRCC5 XRCC6 PRKDC LIG4 MLH1 MSH2 MSH6 FANCD2 RPA1
TP53 MDM2 MDM4 BAX BAK1 BCL2 BCL2L1 MCL1 BBC3 PMAIP1 CASP3 CASP8 CASP9 APAF1 CYCS
FAS XIAP BID
EGFR ERBB2 GRB2 SOS1 HRAS KRAS RAF1 BRAF MAP2K1 MAPK1 MAPK3 PIK3CA AKT1 PTEN MTOR
TSC2 GSK3B CTNNB1 MYC JUN FOS STAT3 JAK2 SRC NFKB1 RELA
""".split()
IMMUNE = """
TLR2 TLR3 TLR4 TLR7 TLR9 MYD88 TICAM1 TIRAP IRAK1 IRAK4 TRAF3 TRAF6 TAB1 TAB2 MAP3K7
IKBKB IKBKG CHUK NFKBIA NFKB1 NFKB2 RELA RELB REL TBK1 IKBKE IRF3 IRF7 IRF1 IRF5
DDX58 IFIH1 MAVS STING1 CGAS IFNA1 IFNB1 IFNG IFNAR1 IFNAR2 IFNGR1 JAK1 JAK2 TYK2
STAT1 STAT2 STAT3 STAT5A STAT6 IRF9 SOCS1 SOCS3 IL1B IL1R1 IL6 IL6R IL6ST IL10 IL10RA
IL2 IL2RA IL4 IL4R TNF TNFRSF1A TRADD RIPK1 RIPK3 CASP8 FADD NLRP3 PYCARD CASP1
MAPK8 MAPK14 JUN FOS CXCL8 CCL2 CD40 CD40LG
""".split()
NETWORKS = {"alignment_network": GENES, "alignment_network_immune": IMMUNE}

# (label, scope, alignment, strength); soft at strength 0 is the independent
# layouts turned rigidly onto the aligned one, so their shapes do not change
STRENGTHS = (0.25, 0.5, 0.75, 1.0)
MODES = (
    [("Per layer (as drawn)", "perLayer", "none", 0.0), ("Soft 0", "perLayer", "soft", 0.0)]
    + [(f"Soft {s:g}", "perLayer", "soft", s) for s in STRENGTHS]
    + [(f"Anchor {s:g}", "perLayer", "anchor", s) for s in STRENGTHS]
    + [(f"Chain {s:g}", "perLayer", "chain", s) for s in STRENGTHS]
    + [("Exact (coupled)", "coupled", "none", 0.0)]
)


def post(path, body):
    req = urllib.request.Request(
        API + path, data=json.dumps(body).encode(), headers={"Content-Type": "application/json"}
    )
    with urllib.request.urlopen(req, timeout=600) as r:
        return json.loads(r.read())


def network(name, genes):
    path = DATA / f"{name}.json"
    if not path.exists():
        net = post("/api/import/databases", {
            "identifiers": genes, "species": 9606,
            "databases": ["STRING", "IntAct", "OmniPath"],
            "required_score": 0.4, "only_among": True,
        })
        path.write_text(json.dumps(net))
    return json.loads(path.read_text())


def layer_graphs(net):
    """Each layer's own graph, over the node ids of that layer."""
    out = {}
    for layer in net["layers"]:
        ids = [n["id"] for n in net["nodes"] if n["layer"] == layer]
        index = {n: i for i, n in enumerate(ids)}
        pairs = {
            tuple(sorted((index[e["src"]], index[e["trg"]])))
            for e in net["edges"]
            if e["source_layer"] == e["target_layer"] == layer and e["src"] != e["trg"]
        }
        out[layer] = (ids, ig.Graph(n=len(ids), edges=sorted(pairs)))
    return out


def structure(positions, graphs):
    """Mean over layers of the Spearman correlation between drawn distances and
    shortest-path distances, over the pairs of each layer's largest component."""
    rhos = []
    for ids, gr in graphs.values():
        comp = max(gr.connected_components(), key=len)
        sp = gr.distances(source=comp, target=comp)
        drawn, graph = [], []
        for a, b in combinations(range(len(comp)), 2):
            pa, pb = positions[ids[comp[a]]], positions[ids[comp[b]]]
            drawn.append(((pa[0] - pb[0]) ** 2 + (pa[1] - pb[1]) ** 2) ** 0.5)
            graph.append(sp[a][b])
        rhos.append(spearmanr(drawn, graph).statistic)
    return statistics.mean(rhos)


def bench(name, genes):
    net = network(name, genes)
    graphs = layer_graphs(net)
    layers = net["layers"]
    inter = sum(1 for e in net["edges"] if e["source_layer"] != e["target_layer"])
    print(f"{len(net['nodes'])} nodes, {len(net['edges'])} edges ({inter} between layers) in {layers}")
    for layer, (ids, gr) in graphs.items():
        print(f"  {layer}: {gr.vcount()} proteins, {gr.ecount()} interactions, largest component {max(len(c) for c in gr.connected_components())}")
    nodes = [NodeModel(id=n["id"], label=n["label"], layer=n["layer"]) for n in net["nodes"]]
    edges = [EdgeModel(**e) for e in net["edges"]]
    rows, same = [], []
    for label, scope, alignment, strength in MODES:
        for seed in SEEDS:
            body = {
                "nodes": net["nodes"], "edges": net["edges"], "algorithm": "Fruchterman-Reingold",
                "scope": scope, "selected_layers": layers, "seed": seed,
                "alignment": alignment, "alignment_strength": strength, "reference_layer": layers[0],
            }
            res = post("/api/layout", body)
            rows.append({"mode": label, "seed": seed, "misalignment": res["misalignment"],
                         "structure": round(structure(res["positions"], graphs), 4)})
            if label == "Per layer (as drawn)":
                extent = max(len(ids) for ids, _ in graphs.values()) ** 0.5
                pos = {}
                for ids, _ in graphs.values():
                    pos.update(_fit({i: tuple(res["positions"][i][:2]) for i in ids}, extent))
                same.append({"mode": "Independent, same size", "seed": seed,
                             "misalignment": misalignment(pos, nodes, edges, layers),
                             "structure": round(structure(pos, graphs), 4)})
        ms = [r["misalignment"] for r in rows if r["mode"] == label]
        st = [r["structure"] for r in rows if r["mode"] == label]
        print(f"{label:22s} misalignment {statistics.mean(ms):6.2f} ± {statistics.stdev(ms):5.2f}   "
              f"structure {statistics.mean(st):.3f} ± {statistics.stdev(st):.3f}")
    rows = same + rows
    ms, st = [r["misalignment"] for r in same], [r["structure"] for r in same]
    print(f"{'Independent, same size':22s} misalignment {statistics.mean(ms):6.2f} ± {statistics.stdev(ms):5.2f}   "
          f"structure {statistics.mean(st):.3f} ± {statistics.stdev(st):.3f}")
    out = DATA / f"{name.replace('network', 'benchmark')}.csv"
    with out.open("w", newline="") as f:
        w = csv.DictWriter(f, fieldnames=list(rows[0]))
        w.writeheader()
        w.writerows(rows)
    print(f"-> {out}\n")


def main():
    for name, genes in NETWORKS.items():
        bench(name, genes)


if __name__ == "__main__":
    main()
```
