## Supplementary material for "Arena3D 3.0: a web server for exploring, analyzing, annotating and sharing multilayered networks in 3D": alignment_panel.py

```
"""Draft figure panel for the alignment benchmark (alignment_benchmark.py).

Left, the STRING, IntAct and OmniPath network as Arena3D 3.0 draws it with
independent layouts (every layer the same size), with soft alignment at
strength 0.5 and with exact alignment. Right, for that network and for the
innate immunity network, the trade-off between misalignment and how well each
layer keeps its own structure, for soft, anchor and chain alignment at
strengths 0 or 0.25 to 1. Means of ten seeds, bars one standard deviation.

With --figure, it instead writes the two charts at their cell size in
Figure 5 (figures/nar_raw/n5d_*.png, n5e_*.png) and copies the three views
there, for compose_nar.py 5.

Run: uv run --no-project --with matplotlib --with pillow --with numpy python alignment_panel.py <views_dir> [--figure]
"""

import csv
import statistics
import sys
from collections import defaultdict
from pathlib import Path

import matplotlib

matplotlib.use("Agg")
import matplotlib.pyplot as plt
import numpy as np
from PIL import Image

HERE = Path(__file__).parent
VIEWS = Path(sys.argv[1]) if len(sys.argv) > 1 else HERE
OUT = HERE / "figures" / "nar_alignment_panel"

# dark-surface categorical slots 1 to 3 (validated all-pairs on black)
SOFT, ANCHOR, CHAIN = "#3987e5", "#d95926", "#199e70"
INK, INK2, GRID, NEUTRAL = "#f0efec", "#c3c2b7", "#2e2e2c", "#e8e7e2"
plt.rcParams.update({"font.family": "Arial", "font.size": 12, "axes.edgecolor": "#5a5955",
                     "axes.labelcolor": INK2, "xtick.color": INK2, "ytick.color": INK2})


def summary(path):
    by = defaultdict(lambda: ([], []))
    for r in csv.DictReader(open(path)):
        by[r["mode"]][0].append(float(r["misalignment"]))
        by[r["mode"]][1].append(float(r["structure"]))
    return {m: (statistics.mean(a), statistics.stdev(a), statistics.mean(b), statistics.stdev(b))
            for m, (a, b) in by.items()}


def view(name):
    """A capture cropped to the network, with a margin."""
    im = np.asarray(Image.open(VIEWS / f"{name}_0.3_-0.7.png").convert("RGB"))
    ys, xs = np.where(im.max(axis=2) > 28)
    pad = 30
    return im[max(ys.min() - pad, 0): ys.max() + pad, max(xs.min() - pad, 0): xs.max() + pad]


def series(ax, s, names, color, label, dashed=False):
    pts = [s[n] for n in names]
    x, xe, y, ye = (np.array(v) for v in zip(*pts))
    ax.errorbar(x, y, xerr=xe, yerr=ye, fmt="none", ecolor=color, elinewidth=1, alpha=0.55, capsize=0)
    ax.plot(x, y, color=color, lw=2, ls=(0, (5, 3)) if dashed else "-", label=label, zorder=3)
    ax.scatter(x, y, s=46, zorder=4, edgecolors=color, linewidths=2,
               facecolors="black" if dashed else color)
    return x, y


def chart(ax, path, title):
    s = summary(path)
    st = ["0.25", "0.5", "0.75", "1"]
    x, y = series(ax, s, ["Soft 0"] + [f"Soft {v}" for v in st], SOFT, "Soft")
    for mode, color, dy in (("Anchor", ANCHOR, -17), ("Chain", CHAIN, 9)):
        xs, ys = series(ax, s, [f"{mode} {v}" for v in st], color, mode)
        # their strengths too (1 sits on the exact layout): anchor below, chain
        # above, on black where a line passes behind
        for xi, yi, v in zip(xs[:-1], ys[:-1], st[:-1]):
            ax.annotate(v, (xi, yi), xytext=(0, dy), textcoords="offset points", ha="center", color=color,
                        fontsize=9, zorder=6, bbox={"facecolor": "black", "edgecolor": "none", "pad": 0.6})
    for xi, yi, v in zip(x[:-1], y[:-1], ["0"] + st[:-1]):
        # left of a rising line: the empty side of the chart
        ax.annotate(v, (xi, yi), xytext=(-9, 3), textcoords="offset points", ha="right", color=INK2, fontsize=10)
    ind, ex = s["Independent, same size"], s["Exact (coupled)"]
    ax.scatter([ind[0]], [ind[2]], s=150, marker="o", facecolors="none", edgecolors=NEUTRAL, linewidths=2, zorder=5)
    ax.annotate("Independent", (ind[0], ind[2]), xytext=(10, 12), textcoords="offset points", ha="right", color=INK)
    ax.scatter([ex[0]], [ex[2]], s=110, marker="D", color=NEUTRAL, zorder=5)
    ax.annotate("Exact", (ex[0], ex[2]), xytext=(4, -24), textcoords="offset points", color=INK)
    ax.set_title(title, color=INK, fontsize=13, loc="left", pad=8)
    ax.set_xlim(-1.5, 31)
    ax.set_ylim(0.25, 0.8)
    ax.set_xlabel("Misalignment (% of the layout diagonal)")
    ax.grid(color=GRID, lw=1)
    ax.set_axisbelow(True)
    ax.set_facecolor("black")
    for side in ("top", "right"):
        ax.spines[side].set_visible(False)


def figure_parts():
    """The charts for Figure 5 at their cell size there (1755 x 680 px, a
    3-column, 2-row cell of compose_nar.py under the letter band), fonts that
    print at about 6.5 pt with the figure 6 to 7 in wide; the views copied."""
    raw = HERE / "figures" / "nar_raw"
    for src, dst in (("soft0", "n5a_independent"), ("soft05fixed", "n5b_soft05"), ("exact", "n5c_exact")):
        if (VIEWS / f"{src}_0.3_-0.7.png").exists():  # else keep the views already in nar_raw
            Image.open(VIEWS / f"{src}_0.3_-0.7.png").save(raw / f"{dst}.png")
    plt.rcParams.update({"font.size": 10.5})
    for key, path, title, legend in (
        ("n5d_tradeoff_dna", HERE / "data" / "alignment_benchmark.csv", "DNA damage and cell cycle", True),
        ("n5e_tradeoff_immune", HERE / "data" / "alignment_benchmark_immune.csv", "Innate immunity", False),
    ):
        fig = plt.figure(figsize=(1755 / 300, 680 / 300), facecolor="black")
        ax = fig.add_axes((0.105, 0.2, 0.875, 0.68))
        chart(ax, path, title)
        ax.set_ylim(0.2, 0.86)
        ax.set_ylabel("Structure kept (ρ)")
        if legend:
            leg = ax.legend(loc="upper left", ncol=3, frameon=False, fontsize=9.5, handlelength=2,
                            columnspacing=1.2, borderaxespad=0.2)
            for t in leg.get_texts():
                t.set_color(INK2)
        fig.savefig(raw / f"{key}.png", dpi=300, facecolor="black")
        plt.close(fig)
        print(raw / f"{key}.png")


def main():
    if "--figure" in sys.argv:
        figure_parts()
        return
    fig = plt.figure(figsize=(21, 4.4), facecolor="black")
    grid = fig.add_gridspec(1, 5, width_ratios=[1, 1, 1, 1.05, 1.05], wspace=0.2, left=0.01, right=0.99, top=0.82, bottom=0.17)
    for k, (name, title, sub) in enumerate([
        ("soft0", "Independent layouts", "misalignment 23.8%, structure kept ρ 0.69"),
        ("soft05fixed", "Soft alignment, strength 0.5", "misalignment 10.8%, structure kept ρ 0.57"),
        ("exact", "Exact alignment", "misalignment 0%, structure kept ρ 0.48"),
    ]):
        ax = fig.add_subplot(grid[0, k])
        ax.imshow(view(name))
        ax.set_axis_off()
        ax.set_title(f"{title}\n", color=INK, fontsize=13, loc="left", pad=4)
        ax.text(0, 1.015, sub, transform=ax.transAxes, color=INK2, fontsize=11, va="bottom")
    a = fig.add_subplot(grid[0, 3])
    chart(a, HERE / "data" / "alignment_benchmark.csv", "DNA damage and cell cycle")
    a.set_ylabel("Layer structure kept (Spearman ρ)")
    b = fig.add_subplot(grid[0, 4], sharey=a)
    chart(b, HERE / "data" / "alignment_benchmark_immune.csv", "Innate immunity")
    plt.setp(b.get_yticklabels(), visible=False)
    # one key for both charts, in a row above them
    handles, labels = a.get_legend_handles_labels()
    leg = fig.legend(handles, labels, loc="upper center", bbox_to_anchor=(0.795, 1.0), ncol=3,
                     frameon=False, fontsize=11, handlelength=2.6, columnspacing=1.6)
    for t in leg.get_texts():
        t.set_color(INK2)
    for ext in ("png", "pdf"):
        fig.savefig(f"{OUT}.{ext}", dpi=200, facecolor="black")
    print(f"{OUT}.png")


if __name__ == "__main__":
    main()
```
