## Supplementary material for "Arena3D 3.0: a web server for exploring, analyzing, annotating and sharing multilayered networks in 3D": ari_panel.py

```
"""Figure 2K: community detection against the planted modules as they weaken
(data/rob4_ari_sweep.csv, from rob4_analyses.py ari). Multiplex Leiden and
Leiden on each layer, mean adjusted Rand index of 20 networks per mixing value
with one standard deviation, in the style of the Figure 5 charts.

Run: uv run --no-project --with matplotlib --with pillow python ari_panel.py
"""

import csv
import statistics
from pathlib import Path

import matplotlib

matplotlib.use("Agg")
import matplotlib.pyplot as plt

HERE = Path(__file__).parent
RAW = HERE / "figures" / "nar_raw"

# dark-surface categorical slots 1 to 4 (validated on the adjacent pairlist),
# multiplex Leiden in the neutral ink of the Figure 5 charts
LAYERS = [("Co-expression", "#3987e5"), ("Protein interactions", "#d95926"),
          ("Genetic interactions", "#199e70"), ("Regulation", "#c98500")]
INK, INK2, GRID, NEUTRAL = "#f0efec", "#c3c2b7", "#2e2e2c", "#e8e7e2"
plt.rcParams.update({"font.family": "Arial", "font.size": 10.5, "axes.edgecolor": "#5a5955",
                     "axes.labelcolor": INK2, "xtick.color": INK2, "ytick.color": INK2})


def main():
    rows = [r for r in csv.DictReader(open(HERE / "data" / "rob4_ari_sweep.csv")) if r["mu"] != "figures"]
    mus = sorted({float(r["mu"]) for r in rows})

    def stats(key):
        vals = [[float(r[key]) for r in rows if float(r["mu"]) == m] for m in mus]
        return [statistics.mean(v) for v in vals], [statistics.stdev(v) for v in vals]

    # the cell of Figure 2K: two columns wide, one row high, under the letter band
    fig = plt.figure(figsize=(2350 / 300, 670 / 300), facecolor="black")
    ax = fig.add_axes((0.075, 0.2, 0.66, 0.7))
    for name, color in LAYERS:
        y, sd = stats(name)
        ax.errorbar(mus, y, yerr=sd, fmt="none", ecolor=color, elinewidth=1, alpha=0.5, capsize=0)
        ax.plot(mus, y, color=color, lw=1.6, label=name, zorder=3)
        ax.scatter(mus, y, s=22, color=color, zorder=4)
    y, sd = stats("multiplex")
    ax.errorbar(mus, y, yerr=sd, fmt="none", ecolor=NEUTRAL, elinewidth=1.2, alpha=0.7, capsize=0)
    ax.plot(mus, y, color=NEUTRAL, lw=2.6, label="Multiplex", zorder=5)
    ax.scatter(mus, y, s=40, color=NEUTRAL, zorder=6)
    ax.set_xlim(0.02, 0.83)
    ax.set_ylim(-0.04, 1.06)
    ax.set_xticks([0.1, 0.2, 0.3, 0.4, 0.5, 0.6, 0.7, 0.8])
    ax.set_xlabel("Mixing μ, the share of a gene's links that leave its module")
    ax.set_ylabel("Adjusted Rand index")
    ax.grid(color=GRID, lw=1)
    ax.set_axisbelow(True)
    ax.set_facecolor("black")
    for side in ("top", "right"):
        ax.spines[side].set_visible(False)
    handles, labels = ax.get_legend_handles_labels()
    order = [labels.index("Multiplex")] + [i for i, t in enumerate(labels) if t != "Multiplex"]
    leg = ax.legend([handles[i] for i in order], [labels[i] for i in order], loc="center left", bbox_to_anchor=(1.02, 0.5),
                    frameon=False, fontsize=9.5, handlelength=2, labelspacing=0.9,
                    title="Leiden", title_fontsize=9.5, alignment="left")
    for t in [*leg.get_texts(), leg.get_title()]:
        t.set_color(INK2)
    fig.savefig(RAW / "n2k_ari_sweep.png", dpi=300, facecolor="black")
    print(RAW / "n2k_ari_sweep.png")


if __name__ == "__main__":
    main()
```
