## Supplementary material for "Arena3D 3.0: a web server for exploring, analyzing, annotating and sharing multilayered networks in 3D": degree_fit.py

```
"""Power-law test of the degree sequences of the DNA damage network's layers
(data/rob4_degrees.json, from rob4_analyses.py degree), with the method of
Clauset et al. (2009): the tail from the k_min that minimizes the KS distance,
and likelihood-ratio tests against an exponential and a lognormal tail.

Run: uv run --no-project --with powerlaw --with mpmath python degree_fit.py
"""

import csv
import json
from pathlib import Path

import powerlaw

DATA = Path(__file__).parent / "data"

rows = []
for layer, degrees in json.loads((DATA / "rob4_degrees.json").read_text()).items():
    fit = powerlaw.Fit([d for d in degrees if d > 0], discrete=True, verbose=False)
    row = {"layer": layer, "nodes": len(degrees), "k_min": fit.xmin, "tail": int(sum(d >= fit.xmin for d in degrees)),
           "gamma": round(fit.alpha, 2), "KS": round(fit.D, 3)}
    for other in ("exponential", "lognormal"):
        r, p = fit.distribution_compare("power_law", other, normalized_ratio=True)
        row[f"vs {other} R"], row[f"vs {other} p"] = round(r, 2), round(p, 3)
    rows.append(row)
    print(row)
with open(DATA / "rob4_degree_fit.csv", "w", newline="") as f:
    w = csv.DictWriter(f, fieldnames=list(rows[0]))
    w.writeheader()
    w.writerows(rows)
```
