## Supplementary material for "Arena3D 3.0: a web server for exploring, analyzing, annotating and sharing multilayered networks in 3D": mock_data6.py

```
"""Six-layer mock data for the NAR figures (as mock_data.py, two more layers) (synthetic, reproducible: seed 7).

mock6_multiplex.tsv        a four-layer multiplex network of 90 genes in six
                          pathway modules (gene symbols are real, the links are
                          random): co-expression, protein interactions, genetic
                          interactions and regulation, each with its own density
                          and hubs; copies of a gene in consecutive layers are
                          linked for about 40% of the genes.
mock6_multiplex_nodes.tsv  node attributes: the module as an annotation term and
                          a one-line description, for hulls, fog clouds, pies
                          and the group layouts.

Run: python3 mock_data.py  (writes data/)
"""

import random
from pathlib import Path

OUT = Path(__file__).parent / "data"
rng = random.Random(7)

MODULES = {
    "Cell cycle": "CDK1 CCNB1 CDK2 CCNE1 CCNA2 PLK1 AURKA BUB1 CDC20 MAD2L1 CDK4 CCND1 E2F1 RB1 CDKN1A",
    "MAPK signaling": "MAPK1 MAPK3 MAP2K1 MAP2K2 BRAF RAF1 KRAS HRAS GRB2 SOS1 EGFR ELK1 FOS JUN DUSP6",
    "PI3K-AKT signaling": "PIK3CA PIK3R1 AKT1 AKT2 PTEN PDK1 MTOR RPTOR RICTOR TSC1 TSC2 RHEB FOXO3 GSK3B IRS1",
    "DNA repair": "BRCA1 BRCA2 RAD51 ATM ATR CHEK1 CHEK2 TP53BP1 MDC1 H2AX XRCC1 PARP1 MRE11 NBN RAD50",
    "Apoptosis": "TP53 MDM2 BAX BAK1 BCL2 BCL2L1 MCL1 CASP3 CASP8 CASP9 APAF1 CYCS BID XIAP FAS",
    "Wnt signaling": "CTNNB1 APC AXIN1 AXIN2 GSK3A LRP6 FZD1 DVL1 TCF7L2 LEF1 WNT3A MYC CCND2 SFRP1 DKK1",
}
module_of = {g: m for m, genes in MODULES.items() for g in genes.split()}
genes = list(module_of)

# per layer: share of genes present, link probability inside a module and
# between modules, and hubs that reach further
LAYERS = [
    ("Co-expression", 0.92, 0.42, 0.003, []),
    ("Protein interactions", 0.88, 0.34, 0.004, ["CDK1", "MAPK1", "AKT1", "TP53", "CTNNB1", "ATM"]),
    ("Genetic interactions", 0.80, 0.26, 0.005, ["BRCA1", "KRAS", "PTEN"]),
    ("Regulation", 0.85, 0.20, 0.002, ["E2F1", "MYC", "FOXO3", "TCF7L2", "JUN", "TP53"]),
    ("Phosphorylation", 0.70, 0.30, 0.003, ["CDK1", "AKT1", "MAPK1", "ATM", "GSK3B"]),
    ("Pathway membership", 0.95, 0.38, 0.001, []),
]

rows = []
present = {}
for layer, share, p_in, p_out, hubs in LAYERS:
    nodes = [g for g in genes if g in hubs or rng.random() < share]
    present[layer] = set(nodes)
    for i, a in enumerate(nodes):
        for b in nodes[i + 1 :]:
            same = module_of[a] == module_of[b]
            p = p_in if same else p_out
            if a in hubs or b in hubs:
                p = min(0.9, p * 1.8 + (0.03 if not same else 0))
            if rng.random() < p:
                rows.append((a, layer, b, layer, round(rng.uniform(0.3, 1.0), 2)))

# copies of a gene in consecutive layers, for about 40% of the genes
coupled = [g for g in genes if rng.random() < 0.4]
for (la, *_), (lb, *_) in zip(LAYERS, LAYERS[1:]):
    for g in coupled:
        if g in present[la] and g in present[lb]:
            rows.append((g, la, g, lb, 1.0))

OUT.mkdir(exist_ok=True)
with open(OUT / "mock6_multiplex.tsv", "w") as f:
    f.write("SourceNode\tSourceLayer\tTargetNode\tTargetLayer\tWeight\n")
    for r in rows:
        f.write("\t".join(map(str, r)) + "\n")
with open(OUT / "mock6_multiplex_nodes.tsv", "w") as f:
    f.write("Node\tLayer\tDescription\tAnnotation\n")
    for layer, *_ in LAYERS:
        for g in sorted(present[layer]):
            m = module_of[g]
            f.write(f"{g}\t{layer}\t{g}, {m.lower()} module (mock data)\t{m}\n")
intra = sum(1 for r in rows if r[1] == r[3])
print(f"{len(rows)} edges ({intra} within layers), {sum(len(p) for p in present.values())} nodes")
```
