## Supplementary material for "Arena3D 3.0: a web server for exploring, analyzing, annotating and sharing multilayered networks in 3D": rob4_analyses.py

```
"""Analyses for Rob's round-4 review (Arena3D_3.0_NAR_rob4_review.md, section 4).

ari         multiplex and single-layer Leiden against the six planted modules of
            the synthetic four-layer network while the modules weaken: the
            mixing mu, the expected share of a gene's links that leave its
            module, goes from 0.05 to 0.8 at the same expected degree per layer
robustness  degree-targeted attack (ranks recalculated) and random failure on
            the DNA damage and cell cycle network of the alignment benchmark
degree      the degree sequences of its layers, for degree_fit.py
enrichment  STRING enrichment of the twelve proteins of Figure 3G, and whether
            each is in the two KEGG pathways drawn
lists       the proteins of both benchmark networks: their layers, degrees and
            KEGG pathways, with nodes and edges per layer
runtimes    layouts, metrics, communities and attacks on networks of 1,000,
            5,000 and 10,000 edges, and the segments for bundling (fdeb_time.mts)

Everything runs in process on the backend's own functions, except the STRING
and KEGG queries. Results go to data/rob4_*.

Run: ../backend/.venv/bin/python rob4_analyses.py [name ...]
"""

import ast
import csv
import json
import random
import statistics
import sys
import time
import urllib.parse
import urllib.request
from pathlib import Path

import igraph as ig
import leidenalg

HERE = Path(__file__).parent
DATA = HERE / "data"
sys.path.insert(0, str(HERE.parent / "backend"))
from app.models.layout import ClusteringRequest, LayoutRequest  # noqa: E402
from app.models.network import EdgeModel, NodeModel  # noqa: E402
from app.models.perturbation import AttackRequest  # noqa: E402
from app.models.topology import TopologyRequest  # noqa: E402
from app.services.layouts import LAYOUTS, compute_clusters, compute_layout  # noqa: E402
from app.services.multilayer import multiplex_membership  # noqa: E402
from app.services.perturbation import attack  # noqa: E402
from app.services.topology import compute_topology  # noqa: E402


def constants(path, *names):
    """Module-level constants of another script, without running it."""
    tree = ast.parse(Path(path).read_text())
    out = {}
    for node in tree.body:
        if isinstance(node, ast.Assign) and isinstance(node.targets[0], ast.Name):
            if node.targets[0].id in names:
                v = node.value
                if isinstance(v, ast.Call) and isinstance(v.func, ast.Attribute) and v.func.attr == "split":
                    out[node.targets[0].id] = ast.literal_eval(v.func.value).split()  # """...""".split()
                else:
                    out[node.targets[0].id] = ast.literal_eval(v)
    return [out[n] for n in names]


def write_csv(name, rows):
    with open(DATA / name, "w", newline="") as f:
        w = csv.DictWriter(f, fieldnames=list(rows[0]))
        w.writeheader()
        w.writerows(rows)


def get(url, data=None):
    req = urllib.request.Request(url, data=data, headers={"User-Agent": "Arena3D-paper-analysis"})
    with urllib.request.urlopen(req, timeout=60) as r:
        return r.read().decode()


# ---------------------------------------------------------------- ARI sweep
MODULES, LAYERS = constants(HERE / "mock_data.py", "MODULES", "LAYERS")
MODULE = {g: m for m, genes in MODULES.items() for g in genes.split()}
GENES = list(MODULE)
MIXING = [0.05, 0.1, 0.2, 0.3, 0.4, 0.5, 0.6, 0.7, 0.8]


def synthetic(mu, seed):
    """The generator of mock_data.py, with the share of links between modules
    set to mu at each layer's expected degree."""
    rng = random.Random(seed)
    layers = {}
    for layer, share, p_in, p_out, hubs in LAYERS:
        nodes = [g for g in GENES if g in hubs or rng.random() < share]
        degree = 14 * p_in + 75 * p_out  # partners in and out of the module
        q_in, q_out = (1 - mu) * degree / 14, mu * degree / 75
        pairs = []
        for i, a in enumerate(nodes):
            for b in nodes[i + 1:]:
                same = MODULE[a] == MODULE[b]
                p = q_in if same else q_out
                if a in hubs or b in hubs:
                    p = min(0.9, p * 1.8 + (0.03 if not same else 0))
                if rng.random() < p:
                    pairs.append((a, b))
        layers[layer] = pairs
    return layers


def ari(names, membership):
    truth = [list(MODULES).index(MODULE[n]) for n in names]
    return ig.compare_communities(truth, membership, method="ari")


def scores(layers, seed):
    """ARI of multiplex Leiden (every gene in any layer) and of Leiden on each
    layer (the genes of that layer), with the observed mixing per layer."""
    edges = []
    for layer, pairs in layers.items():
        for a, b in pairs:
            edges.append(EdgeModel(src=f"{a}_{layer}", trg=f"{b}_{layer}", source_node=a, source_layer=layer,
                                   target_node=b, target_layer=layer, weight=1, scaled_weight=1))
    labels = sorted({n for pairs in layers.values() for p in pairs for n in p})
    multi = multiplex_membership(edges, labels, list(layers), seed)
    out = {"multiplex": ari(labels, [multi[n] for n in labels])}
    for layer, pairs in layers.items():
        names = sorted({n for p in pairs for n in p})
        index = {n: i for i, n in enumerate(names)}
        graph = ig.Graph(n=len(names), edges=[(index[a], index[b]) for a, b in pairs])
        part = leidenalg.find_partition(graph, leidenalg.ModularityVertexPartition, seed=seed)
        out[layer] = ari(names, part.membership)
        out[f"{layer} mixing"] = sum(MODULE[a] != MODULE[b] for a, b in pairs) / max(1, len(pairs))
    return out


def run_ari():
    rows = []
    # the synthetic network of the figures (mock_data.py, seed 7)
    net = {layer: [] for layer, *_ in LAYERS}
    with open(DATA / "synthetic_multiplex.tsv") as f:
        for r in csv.DictReader(f, delimiter="\t"):
            if r["SourceLayer"] == r["TargetLayer"]:
                net[r["SourceLayer"]].append((r["SourceNode"], r["TargetNode"]))
    for seed in range(1, 21):
        rows.append({"mu": "figures", "network_seed": 7, "leiden_seed": seed, **scores(net, seed)})
    for mu in MIXING:
        for seed in range(1, 21):
            rows.append({"mu": mu, "network_seed": seed, "leiden_seed": seed, **scores(synthetic(mu, seed), seed)})
    write_csv("rob4_ari_sweep.csv", rows)
    layer_names = [layer for layer, *_ in LAYERS]
    print(f"{'mu':>8} {'multiplex':>9} " + " ".join(f"{n[:12]:>12}" for n in layer_names) + "  mixing")
    for mu in ["figures", *MIXING]:
        sel = [r for r in rows if r["mu"] == mu]
        mean = lambda k: statistics.mean(r[k] for r in sel)  # noqa: E731
        print(f"{mu!s:>8} {mean('multiplex'):9.2f} " + " ".join(f"{mean(n):12.2f}" for n in layer_names)
              + "  " + " ".join(f"{mean(n + ' mixing'):.2f}" for n in layer_names))


# ---------------------------------------------------------------- the DNA damage network
def ddr():
    net = json.loads((DATA / "alignment_network.json").read_text())
    nodes = [NodeModel(id=n["id"], label=n["label"], layer=n["layer"]) for n in net["nodes"]]
    edges = [EdgeModel(**e) for e in net["edges"]]
    return nodes, edges, [layer["name"] if isinstance(layer, dict) else layer for layer in net["layers"]]


def run_robustness():
    nodes, edges, layers = ddr()
    rows = []
    for strategy in ("degree", "random"):
        res = attack(AttackRequest(nodes=nodes, edges=edges, layers=layers, strategy=strategy,
                                   recalculate=True, runs=20, steps=50, seed=1))
        for c in res.curves:
            rows.append({"strategy": strategy, "scope": c.name, "nodes": c.nodes, "R": round(c.robustness, 3),
                         "fraction": json.dumps(c.fraction), "largest": json.dumps(c.largest)})
            print(f"{strategy:7} {c.name:15} n={c.nodes:4} R={c.robustness:.3f}")
    write_csv("rob4_robustness.csv", rows)


def run_degree():
    nodes, edges, layers = ddr()
    out = {}
    for layer in layers:
        ids = [n.id for n in nodes if n.layer == layer]
        index = {v: i for i, v in enumerate(ids)}
        pairs = {tuple(sorted((index[e.src], index[e.trg]))) for e in edges
                 if e.source_layer == layer == e.target_layer and e.src != e.trg}
        out[layer] = ig.Graph(n=len(ids), edges=sorted(pairs)).degree()
    (DATA / "rob4_degrees.json").write_text(json.dumps(out))
    for layer, deg in out.items():
        print(f"{layer:10} n={len(deg)} edges={sum(deg) // 2} max degree={max(deg)}")


# ---------------------------------------------------------------- Figure 3G
FIG3G = "BRCA1 BRCA2 RAD51 PALB2 BARD1 ATM CHEK2 TP53 MDM2 CDKN1A BAX BBC3".split()


def kegg_genes(pathway):
    """Gene symbols of a KEGG pathway (the GENE section of its entry)."""
    genes, section = [], None
    for line in get(f"https://rest.kegg.jp/get/{pathway}").splitlines():
        if line[:12].strip():
            section = line[:12].strip()
        if section == "GENE":
            genes.append(line[12:].split(";")[0].split()[1])
    return genes


def run_enrichment():
    version = json.loads(get("https://string-db.org/api/json/version"))[0]
    body = urllib.parse.urlencode({"identifiers": "\r".join(FIG3G), "species": 9606,
                                   "caller_identity": "Arena3D-paper"}).encode()
    terms = json.loads(get("https://string-db.org/api/json/enrichment", body))
    kegg = sorted((t for t in terms if t["category"] == "KEGG"), key=lambda t: t["fdr"])
    rows = [{"term": t["term"], "description": t["description"], "genes": t["number_of_genes"],
             "fdr": t["fdr"], "proteins": ",".join(t["preferredNames"])} for t in kegg]
    write_csv("rob4_fig3g_kegg.csv", rows)
    print(f"STRING {version.get('string_version')}: {len(kegg)} KEGG pathways at FDR < 0.05")
    for r in rows:
        print(f"  {r['term']} {r['description'][:40]:40} {r['genes']:2} genes FDR {r['fdr']:.1e}")
    for pathway in ("hsa03440", "hsa04115"):
        genes = set(kegg_genes(pathway))
        print(f"{pathway}: {', '.join(g for g in FIG3G if g in genes)} | ATM in it: {'ATM' in genes}")


# ---------------------------------------------------------------- benchmark lists
KEGG_PATHWAYS = {
    "hsa04110": "Cell cycle", "hsa04115": "p53 signaling", "hsa03440": "Homologous recombination",
    "hsa03450": "Non-homologous end joining", "hsa03430": "Mismatch repair", "hsa03460": "Fanconi anemia",
    "hsa04210": "Apoptosis", "hsa04010": "MAPK signaling", "hsa04151": "PI3K-Akt signaling",
    "hsa04012": "ErbB signaling", "hsa04630": "JAK-STAT signaling", "hsa04310": "Wnt signaling",
    "hsa04150": "mTOR signaling", "hsa04064": "NF-kappa B signaling", "hsa04620": "Toll-like receptor signaling",
    "hsa04621": "NOD-like receptor signaling", "hsa04622": "RIG-I-like receptor signaling",
    "hsa04623": "Cytosolic DNA-sensing", "hsa04668": "TNF signaling", "hsa04060": "Cytokine-cytokine receptor",
}


def run_lists():
    genes, immune = constants(HERE / "alignment_benchmark.py", "GENES", "IMMUNE")
    pathways = {}
    for pid, name in KEGG_PATHWAYS.items():
        for g in kegg_genes(pid):
            pathways.setdefault(g, []).append(name)
        time.sleep(0.4)
    rows, summary = [], []
    for title, path, proteins in (("DNA damage and cell cycle", "alignment_network.json", genes),
                                  ("Innate immunity", "alignment_network_immune.json", immune)):
        net = json.loads((DATA / path).read_text())
        layers = [layer["name"] if isinstance(layer, dict) else layer for layer in net["layers"]]
        degree = {}
        inter = 0
        for e in net["edges"]:
            if e["source_layer"] != e["target_layer"]:
                inter += 1
                continue
            for n in (e["src"], e["trg"]):
                degree[n] = degree.get(n, 0) + 1
        for layer in layers:
            n = sum(1 for v in net["nodes"] if v["layer"] == layer)
            m = sum(1 for e in net["edges"] if e["source_layer"] == layer == e["target_layer"])
            summary.append({"network": title, "layer": layer, "nodes": n, "edges": m})
        summary.append({"network": title, "layer": "links between layers", "nodes": "", "edges": inter})
        present = {(v["label"], v["layer"]) for v in net["nodes"]}
        for p in proteins:
            rows.append({"network": title, "protein": p,
                         **{f"{layer} degree": degree.get(f"{p}_{layer}", 0) if (p, layer) in present else ""
                            for layer in layers},
                         "KEGG pathways": "; ".join(pathways.get(p, []))})
        covered = sum(1 for p in proteins if p in pathways)
        print(f"{title}: {len(proteins)} proteins, {covered} in the {len(KEGG_PATHWAYS)} KEGG pathways")
    for s in summary:
        print(f"  {s['network']:26} {s['layer']:22} nodes {s['nodes']!s:>4} edges {s['edges']}")
    with open(DATA / "Table_S1_benchmark_proteins.tsv", "w", newline="") as f:
        fields = sorted({k for r in rows for k in r}, key=lambda k: list(rows[0]).index(k) if k in rows[0] else 99)
        w = csv.DictWriter(f, fieldnames=fields, delimiter="\t")
        w.writeheader()
        w.writerows(rows)
    write_csv("rob4_benchmark_layers.csv", summary)


# ---------------------------------------------------------------- runtimes
SIZES = [1000, 5000, 10000]
N_LAYERS = 4


def random_network(n_edges, seed=1):
    """Four Barabasi-Albert layers (two links per new node) over shared names,
    n_edges within layers in total."""
    per_layer = n_edges // N_LAYERS
    n = per_layer // 2 + 1
    nodes, edges = [], []
    for k in range(N_LAYERS):
        layer = f"Layer{k + 1}"
        g = ig.Graph.Barabasi(n, 2)
        random.Random(seed + k).shuffle(names := [f"G{i}" for i in range(n)])
        for v in range(n):
            nodes.append(NodeModel(id=f"{names[v]}_{layer}", label=names[v], layer=layer))
        for a, b in g.get_edgelist()[:per_layer]:
            edges.append(EdgeModel(src=f"{names[a]}_{layer}", trg=f"{names[b]}_{layer}", source_node=names[a],
                                   source_layer=layer, target_node=names[b], target_layer=layer,
                                   weight=1, scaled_weight=1))
    return nodes, edges, [f"Layer{k + 1}" for k in range(N_LAYERS)]


def timed(fn, repeat=1):
    best = None
    for _ in range(repeat):
        t = time.perf_counter()
        out = fn()
        dt = time.perf_counter() - t
        best = dt if best is None else min(best, dt)
    return best, out


def run_runtimes():
    rows = []
    # UMAP compiles its numba code on the first call: not part of a run time
    nodes, edges, layers = random_network(200)
    compute_layout(LayoutRequest(nodes=nodes, edges=edges, algorithm="UMAP", scope="perLayer", selected_layers=layers))
    for size in SIZES:
        nodes, edges, layers = random_network(size)
        jobs = {}
        for algo in [*LAYOUTS, "UMAP"]:
            jobs[f"layout: {algo}"] = lambda a=algo: compute_layout(
                LayoutRequest(nodes=nodes, edges=edges, algorithm=a, scope="perLayer", selected_layers=layers))
        for mode, strength in (("soft", 0.5), ("anchor", 0.5), ("chain", 0.5)):
            jobs[f"alignment: {mode} 0.5"] = lambda m=mode, s=strength: compute_layout(
                LayoutRequest(nodes=nodes, edges=edges, algorithm="Fruchterman-Reingold", scope="perLayer",
                              selected_layers=layers, alignment=m, alignment_strength=s))
        jobs["alignment: exact"] = lambda: compute_layout(
            LayoutRequest(nodes=nodes, edges=edges, algorithm="Fruchterman-Reingold", scope="coupled",
                          selected_layers=layers))
        for metric in ("Degree", "Clustering Coefficient", "Betweenness Centrality", "Closeness Centrality",
                       "Eigenvector Centrality", "PageRank", "Coreness (k-core)", "Eccentricity"):
            jobs[f"metric: {metric}"] = lambda m=metric: compute_topology(
                TopologyRequest(nodes=nodes, edges=edges, metric=m, scope="perLayer", selected_layers=layers))
        for algo in ("Louvain", "Walktrap", "Fast Greedy", "Label Propagation", "Leiden"):
            jobs[f"communities: {algo}"] = lambda a=algo: compute_clusters(
                ClusteringRequest(nodes=nodes, edges=edges, algorithm=a, scope="perLayer", selected_layers=layers))
        jobs["communities: multiplex Leiden"] = lambda: compute_clusters(
            ClusteringRequest(nodes=nodes, edges=edges, algorithm="Leiden", scope="coupled", selected_layers=layers))
        jobs["attack: degree, recalculated"] = lambda: attack(
            AttackRequest(nodes=nodes, edges=edges, layers=layers, strategy="degree", recalculate=True))
        jobs["attack: random failure, 5 runs"] = lambda: attack(
            AttackRequest(nodes=nodes, edges=edges, layers=layers, strategy="random", runs=5))
        for name, fn in jobs.items():
            try:
                seconds, _ = timed(fn)
                rows.append({"edges": size, "nodes": len(nodes), "task": name, "seconds": round(seconds, 3)})
            except Exception as exc:  # a layout the server refuses at this size
                rows.append({"edges": size, "nodes": len(nodes), "task": name, "seconds": f"refused: {exc}"[:120]})
            print(f"{size:6} {name:40} {rows[-1]['seconds']}", flush=True)
        # the segments the client bundles: each layer's edges on its floor (FR)
        pos, _ = compute_layout(LayoutRequest(nodes=nodes, edges=edges, algorithm="Fruchterman-Reingold",
                                              scope="perLayer", selected_layers=layers))
        segments = {}
        for e in edges:
            (ay, az), (by, bz) = pos[e.src][:2], pos[e.trg][:2]
            x = layers.index(e.source_layer) * 100.0
            segments.setdefault(e.source_layer, []).append([[x, ay * 100, az * 100], [x, by * 100, bz * 100]])
        (DATA / f"rob4_segments_{size}.json").write_text(json.dumps(segments))
    write_csv("rob4_runtimes.csv", rows)


RUNS = {"ari": run_ari, "robustness": run_robustness, "degree": run_degree, "enrichment": run_enrichment,
        "lists": run_lists, "runtimes": run_runtimes}

if __name__ == "__main__":
    for name in sys.argv[1:] or list(RUNS):
        print(f"== {name}", flush=True)
        RUNS[name]()
```
